# Environmental enrichment induces epigenomic and genome organization changes relevant for cognitive function

**DOI:** 10.1101/2021.01.31.428988

**Authors:** Sergio Espeso-Gil, Aliaksei Holik, Sarah Bonnin, Shalu Jhanwar, Sandhya Chandrasekaran, Roger Pique-Regi, Júlia Albaigès-Ràfols, Michael Maher, Jon Permanyer, Manuel Irimia, Marc R. Friedländer, Meritxel Pons-Espinal, Schahram Akbarian, Mara Dierssen, Philipp G. Maass, Charlotte N. Hor, Stephan Ossowski

## Abstract

In early development, the environment triggers mnemonic epigenomic programs resulting in memory and learning experiences to confer cognitive phenotypes into adulthood. To uncover how environmental stimulation impacts the epigenome and genome organization, we used the paradigm of environmental enrichment (EE) in young mice constantly receiving novel stimulation. We profiled epigenome and chromatin architecture in whole cortex and sorted neurons by deep-sequencing techniques. Specifically, we studied chromatin accessibility, gene and protein regulation, and 3D genome conformation, combined with predicted enhancer and chromatin interactions. We identified increased chromatin accessibility, transcription factor binding including CTCF-mediated insulation, differential occupancy of H3K36me3 and H3K79me2, and changes in transcriptional programs required for neuronal development. EE stimuli led to local genome re-organization by inducing increased contacts between chromosomes 7 and 17 (*inter-*chromosomal). Our findings support the notion that EE-induced learning and memory processes are directly associated with the epigenome and genome organization.

**Highlights:**

- Environmental enrichment (EE) alters chromatin conformation, CTCF binding, and spatially 3D genome changes, thereby regulating cognitive function during the first steps of life after birth.
- Transcription-associated gene body marks H3K79me2 and H3K36me3 are differently influenced by EE in cortical brain cells and binding is exacerbated upon stimulation in an age-dependent manner.
- EE-induced changes of 3D genome organization increase *inter-*chromosomal interactions of genes associated with synaptic transmission and AMPA receptor genes on chromosomes 7 and 17.

## Introduction

Exposure to environmental stimuli influences developmental programs of organisms by modulating gene regulatory networks. These programs direct early postnatal neuronal development, particularly during the “critical period” that is key to establish brain functions that are kept throughout the lifetime of an individual (Hübener and Bonhoeffer, 2014). Environmental enrichment (EE) is a commonly used paradigm to study the behavioral and electrophysiological mechanisms of neuronal development (van Praag et al., 2000). EE represents external factors, such as sensory, physical, cognitive, and social stimulation to provide and to maintain the brain with constant novelty and complexity, thereby laying a key foundation for future learning processes (Rountree-Harrison et al., 2018).

The coalescing mechanisms of gene regulation, epigenetics, and genome organization leading to learning and memory formation still remain largely unknown. Thus far, studies on how EE affects gene regulatory elements are sparse, but some findings point towards the involvement of epigenetic mechanisms, both at the level of DNA methylation and histone modifications and chromatin modifiers (Irier et al., 2014; Morse et al., 2015). Recently, advances in brain research indicated that three-dimensional (3D) genome organization can be causally involved in gene-regulatory networks and chromatin conformation dynamics that impact brain function, learning, and memory formation (Rajarajan et al., 2019). These findings imply that a comprehensive molecular analysis of the processes happening during EE is needed to understand how neuronal circuits are refined by environmental cues. To accomplish this aim, we leveraged multiple genomic techniques to identify regulatory changes leading to learning and memory formation by EE during early postnatal neuronal development. We assessed chromatin accessibility, chromatin modifications, transcriptomic and proteomic changes, as well as 3D genome conformation. Our results reveal for the first time a comprehensive genome-wide perspective of global and neuronal-specific regulatory epigenetic modifications under EE in whole cerebral cortex, followed by neuron-specific and pyramidal-neuron-specific profiling. Our present study demonstrates that EE-induced early learning experience changes the neuronal epigenome and causes altered genomic conformations, especially between different chromosomes.

## Results

### Study outline

EE significantly influences learning and memory and leads to cognitive improvement, as demonstrated by multiple studies (Ohline and Abraham, 2019). Our EE-protocol was successfully established and validated by behavioral testing (Morris water maze) in an earlier study (Pons-Espinal et al., 2013). Briefly it consisted in constantly-changing cognitive stimulation (every 48h) over the course of one week (postnatal day P28) and a month (P51), which reflect important stages of the critical period (Hübener and Bonhoeffer, 2014; Sztainberg and Chen, 2010)(Method Details, Figure 1A). For detailed insights into cell heterogeneity in the context of EE and the cerebral cortex microenvironment, we used whole cerebral cortex, sorted neuronal, and non-neuronal cells and performed epigenomic, transcriptomic, and proteomic profiling, as well as capturing of genomic interactions to provide a communal resource (Figure 1B, C).

**Figure1.**
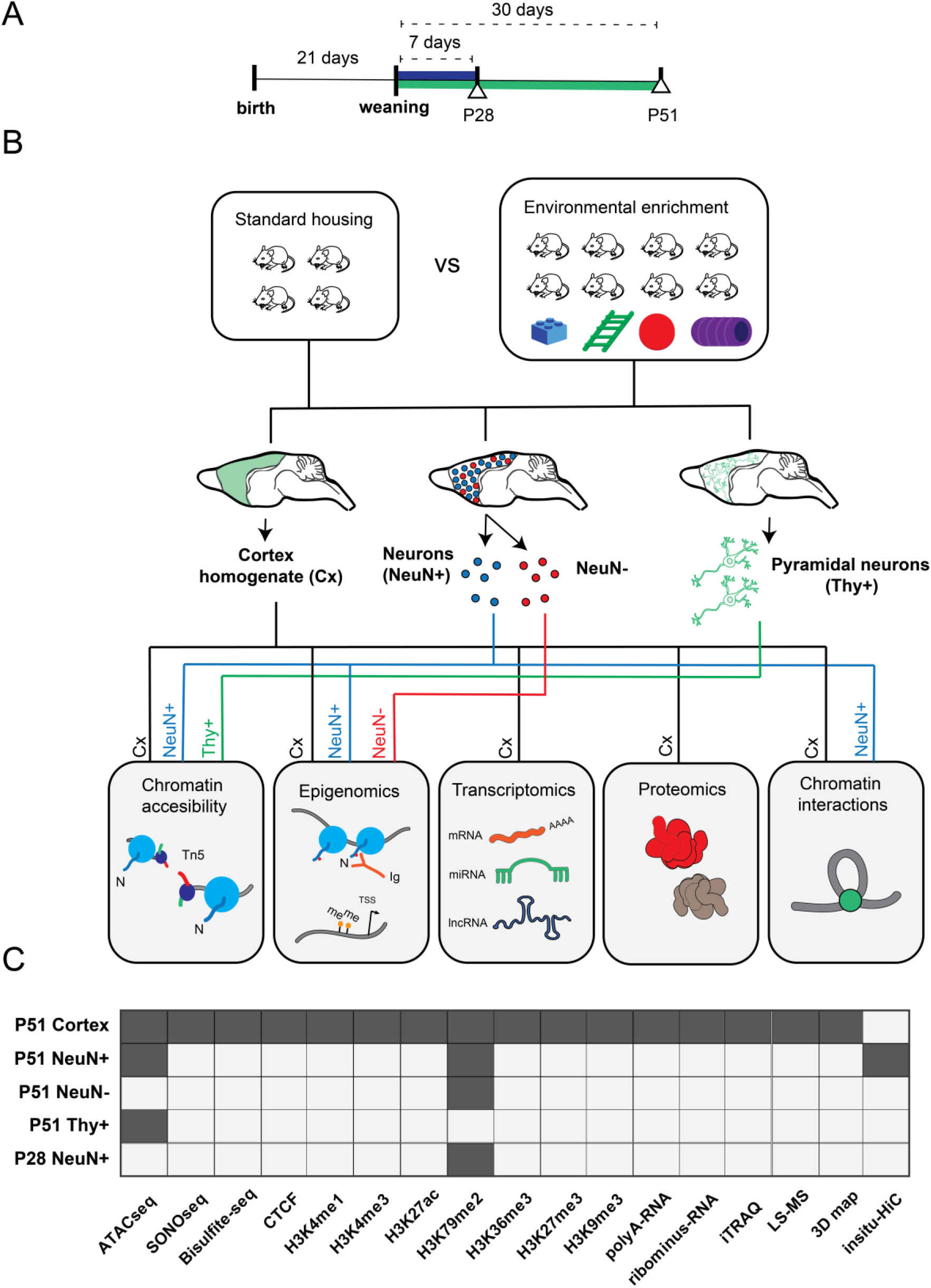
Experimental study design. **A)** After weaning (P21), mice were exposed to environmental enrichment (EE) for 7 days (P28), and 30 days (P51, Methods). **B)** Experimental workflow. Cortical tissue was homogenized from 5 different animals and split for the following protocols: ATACseq/SONOseq, ChIP-seq, RNAseq, label free and iTRAQ proteomics, and *in situ* Hi-C (2 biological replicates per condition, *N*_*t*_=20 animals in total). Neuronal and glial populations were sorted by the neuronal marker NeuN (Rbfox3) and pyramidal neurons by Thy+ (Tg[Thy1-YFP] mice). NeuN^+^ and NeuN^-^ (3 biological replicates per condition, *N*_*t*_=30 of animals; for Thy+ 2 individual biological replicates per condition, *N*_*t*_=4 animals; see Methods. **C)** Datasets available per technique and per different cell population (dark grey).

To analyze and intersect our multiple datasets, we devised a computational pipeline to determine activity and interplay of epigenomic marks in gene-regulatory regions, namely between enhancers predicted by the tool GEP (Jhanwar et al., 2018), and annotated promoters (Methods Details, Figure 2A, Table S1).

**Figure 2.**
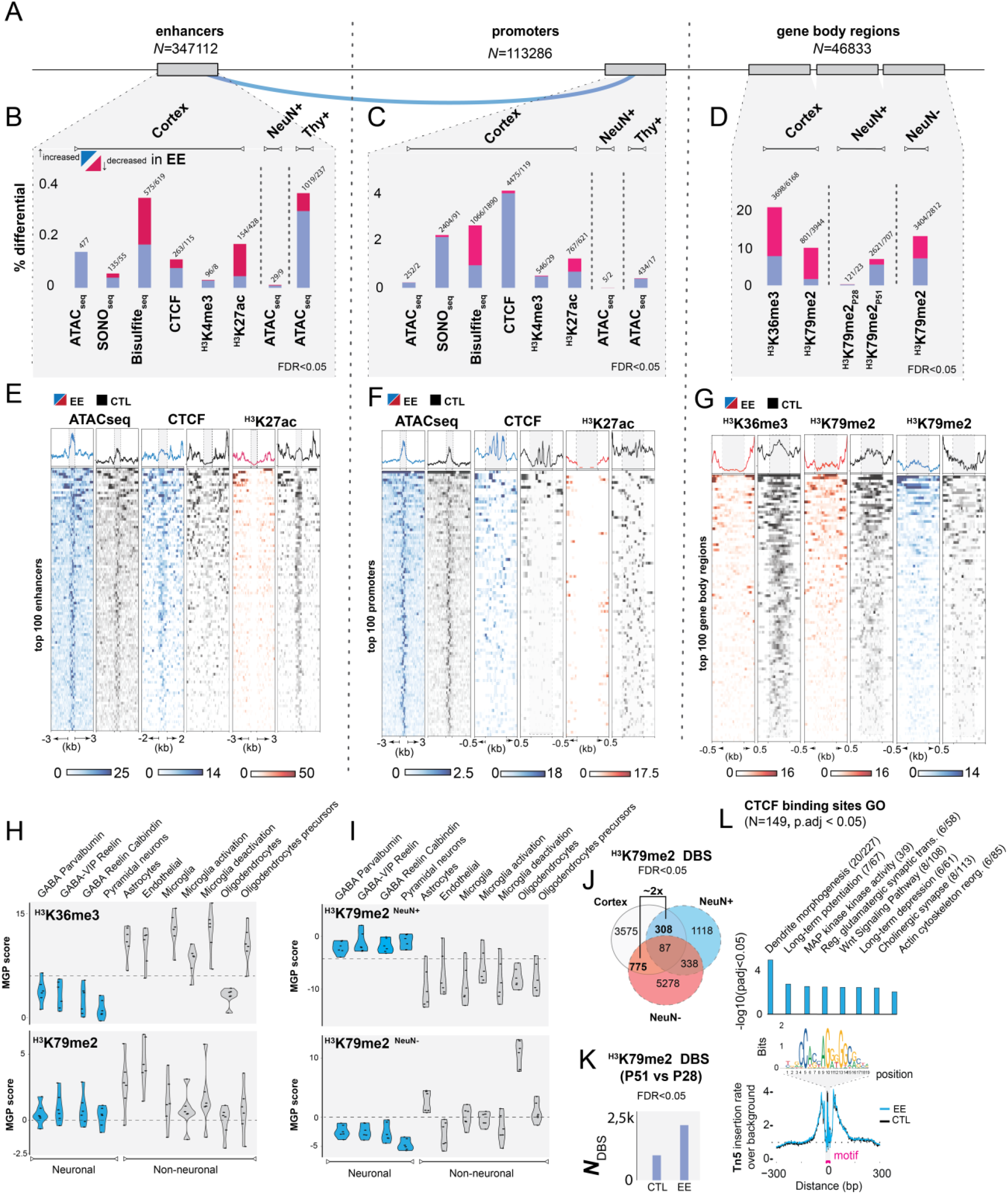
EE epigenetic changes during postnatal development. **A)**Genomic features studied in the present study: left, enhancers predicted by GEP (Jhanwar et al., 2018) (*N*_*t*_=347112), middle: promoters 1500 bp up- and 500 bp down-stream of TSS (*N*_*t*_=113286); right: gene body regions (*N*_*t*_=46833; see Methods). **B-D)** Summary of differential changes [%] upon EE of chromatin accessibility and epigenetic marks over the total number of features in B) enhancers, C) promoters, D) and gene-body regions (FDR<0.05). **E)** Top 100 enhancers, **F)** promoters, and **G)** Gene-body regions scaled in RPKM of the most important marks. Blue = increased; red = decreased signal upon EE, black = CTL samples. **H**,**I)** Cell deconvolution of transcription-associated gene body marks: H3K36me3 and H3K79me2 in both whole cortex, neuronal and non-neuronal datasets. Marker gene profile score (MPG) represents the first principal component regarding gene expression of cell-specific genes curated from single-cell studies involving GABAergic and pyramidal neurons, astrocytes, oligodendrocytes, microglia and endothelial cells (Mancarci et al., 2017) (Methods Details). **J)** Overlap of differential H3K79me2 enrichment at P51 of whole cortex with NeuN^+^ and NeuN^-^ (at FDR<0.05). **K)** Time-course plot showing the progressive increase of differential binding sites (DBS) of H3K79me2 (P51 vs P28) in CTL and EE samples (FDR<0.05). **L**) NeuN^+^ CTCF footprint plot. Y-axis corresponds to the Tn5 insertion rate over the background, x-axis distance in bp from the motif center (upper plot: bins over nucleotide position). Blue line designates increased CTCF binding in EE samples. Right plot: GO analysis (p-adj <0.05 with Benjamini-Holchberg correction, Table S2)

### EE induces increased chromatin accessibility and insulation targeting synaptic-associated genes in cortical tissue

EE is non-invasive in comparison to invasive neuronal stimulation which leads to increased chromatin accessibility in gene-regulatory regions to induce gene transcription (Fernandez-Albert et al., 2019; Koberstein et al., 2018a; Su et al., 2017). Therefore, we asked if non-invasive EE could lead to quantifiable effects on gene and genome function during cortical cell postnatal development.

First, we studied EE in whole cortical tissue (Pons-Espinal et al., 2013). In ATAC-seq experiments investigating chromatin accessibility (Buenrostro et al., 2013), we observed that distinct ATAC-seq peaks (macs2, fseq) were increased genome-wide in EE samples compared to controls (CTLs), suggesting a global increase in chromatin accessibility after EE (FC_cortex_1.17X, Figure S1A). To validate these findings, a differential analysis of enhancers and promoters further confirmed increased chromatin accessibility in a very specific set of 0.13% of enhancers and 0.22 % of the total promoters (FDR<0.05, Figures 2B-C). To link *intra-*chromosomal interactions of enhancers to their corresponding promoters, we used a modified version of EpiTensor (Zhu et al., 2016)(Table S1, Method details). We found regulatory regions showing increased accessibility specific to genes that could be linked to neurogenesis and differentiation (Clemenson et al., 2015; Speisman et al., 2013), angiogenesis (Yu et al., 2014), synapse organization (Ohline and Abraham, 2019), and pathways associated to memory and learning such as Wnt (He et al., 2018), and Rho signaling (Martino et al., 2013) (Figures S1B,C, Table S2). To further confirm previous ATAC-seq results, we used SONO-seq (Auerbach et al., 2009), a method based solely on sonicated and sequenced chromatin. We validated 76 genes showing consistent increased accessibility in their enhancers and promoters (p-adj<0.01, Figure S1E). Additionally, SONO-seq showed differential accessibility on pathways that are important in neuronal function such as MAPK and JNK (Coffey, 2014), neural maturation BMP (Bond et al., 2012), synaptic plasticity PI3K-Akt (Tan et al., 2017), cellular aging prevention and telomere protective role of oxytocin (Faraji et al., 2018; Stevenson et al., 2019), and neurotransmission function by GPCR (Betke et al., 2012) (Table S2).

Accessible regions of chromatin regions encompass characteristic posttranslational modifications in surrounding histones (Fu et al., 2018). Due to the relationship between these histone marks and the increased chromatin accessibility in enhancers and promoters upon EE, we hypothesized that EE could also modulate posttranslational histone modifications and CTCF binding in gene-regulatory regions. We investigated a variety of histone marks from active (H3K27ac, H3K4me3, H3K4me1) to repressed regions (H3K27me3, H3K9me3), in addition to CTCF and DNA methylation. Interestingly, we detected relevant changes in H3K4me3, H3K27ac, DNA methylation and CTCF upon EE (Table S3). The regions represented about 0.2 -0.4% of enhancers and 2 -5% of the promoters depending on the mark being analyzed (FDR<0.05, Figures 2B, C, E-F, Table S3). With the exception of hypermethylated sites and weak changes in H3K27me3, the majority of changes occurred in activity-associated marks. We did not find significant changes in the heterochromatic mark H3K9me3, indicating that EE impacts the modulation of active gene sets rather than repressed regions. To investigate this in detail, we profiled transcription-associated marks such as H3K36me3 and H3K79me2 as potential readouts of gene expression (Huff et al., 2010), and determined a ∼20% and 10% differential binding of gene body marks respectively, confirming that EE impacts transcriptional programs (FDR<0.05, Figures 2D,G, Table S3). Remarkably, the modulation of transcription-associated marks post EE induction was also observed in ∼13% and 8% of distal enhancers bearing H3K36me3 and H3K79me2 respectively, suggesting that enhancer-derived RNA genes are also differentially expressed (Kim et al., 2010) (FDR<0.05, Figures S1F,G). Genes associated to EE-induced cortical epigenetic marks changes were linked to the extracellular matrix (ECM) important for shaping synapses during postnatal development (Bikbaev et al., 2015; Ferrer-Ferrer and Dityatev, 2018), to circadian clock genes known to be required for proper healthy adult behavior (Brooks and Canal, 2013), and glutamatergic receptor complexes key for neuronal plasticity (Lüscher and Malenka, 2012) (Figures S1H-N, Table S3).

Having determined that EE induced differential chromatin accessibility and modulation of histone modifications in postnatal cortical tissue, we next addressed potential cross-talk mechanisms. We explored the overlap across all differential epigenetically modified and accessible chromatin regions identified previously (Figure S1O). The strongest overlap corresponded to increased CTCF binding (18.7% of total sites) co-occurring with a decrease of the gene body activity marks H3K79me2 and H3K36me3 (at FDR<0.1, Figure S1P). A highly relevant example of this priming state is the early-life stress gene *Met* (Heun-johnson and Levitt, 2018), and the memory regulating phosphodiesterase *Pde8b* (Tsai et al., 2012), both showing increased chromatin accessibility of interacting enhancers upon EE, as well as increased CTCF insulation in promoters, but decreased occupancy of H3K79me2 and H3K36me3 when compared to CTLs (Figures S2A,B). This result could indicate a state where genes are poised to be transcribed, but are temporarily repressed by insulation, a specific state due to changes in chromatin architecture (Kim et al., 2015). But, it could also indicate mixed signals coming from the process of synaptic tuning, where some synapses gain strength meanwhile others are lost as consequence of the learning mechanism (Turrigiano, 2008).

### Molecular phenotypic changes by EE mainly target the glutamatergic synapse

Next, we determined how the previously described epigenetic changes alter transcriptional (coding and non-coding) and translational landscapes upon EE. Expression analysis revealed a total of 473 differentially expressed genes (FDR<0.05, Figure S2C, Table S4). Additional biological replicates and gene ontology analysis recapitulated previously described pathways and terms, such as: BMP, JNK, MAPK, AMPAR signaling, and elements of the ECM (Figures S2D-F). By investigating the non-coding fraction of RNA, we identified 200 microRNAs and 52 long non-coding RNAs (lncRNAs) differentially expressed upon EE (Figure S2G-J, Table S5). Top microRNAs were validated in a new set of biological replicates (Figure S2H). Using a multi-source microRNA target predictor (Friedman et al., 2010), we observed specificity for synaptic-associated mRNA targets in a gene ontology analysis (p-adj<0.05, Figure S2I,J, Table S5). Of note, we found *Meg3* and *Rian* (*Meg8*) as downregulated lncRNAs upon EE. Both are known for their ability to regulate glutamatergic neurotransmission, potentially in collaboration with microRNAs (Tan et al., 2017). We then explored the potential interactions between both non-coding elements (microRNAs and lnRNAs) with LncBase (Paraskevopoulou et al., 2016), and observed that 20 of our differentially expressed microRNAs could interact with *Meg3* (Figure S2K). Particularly interesting is the up-regulated miR125b-p5 reported to be involved in synaptic strength and *Grin2a* downregulation (Edbauer et al., 2010).

To recapitulate EE-induced changes by quantitative protein expression, we used iTRAQ and LCMS mass spectrometry, finding about 73 and 145 differential proteins respectively (p-val<0.05, Table S6). Gene ontology analysis of intersected EE-induced transcriptomic and proteomic changes identified pathways highlighting the importance of the ECM and neurotransmission receptor complexes, particularly involving the glutamatergic synapse (Figure S2L).

### EE stimulation in NeuN+ sorted neurons

To further understand the poised state of genes observed in cortical tissue and to avoid cell bias composition, we decided to investigate EE-induced influence in a cell-specific manner. We performed a cell deconvolution analysis to specify which cell types are primarily responding to EE stimulation (Mancarci et al., 2017). Remarkably, we observed that H3K36me3 and H3K79me2 were enriched in non-neuronal populations in whole cortex data (Figure 2H). However, to address the neuronal extent of EE-induced epigenetic changes observed in whole cortex, we performed FACSorting by nuclei immunostaining of the neuronal specific marker NeuN (Jiang et al., 2008) (Rbfox3, Figure S3A). Another deconvolution of H3K79me2 in sorted populations demonstrated the neuronal-specific identification of EE-stimulatory effects (Figure 2I, Figure S3B). We observed that differential analysis on H3K79me2 data between EE *versus* CTLs showed two times more non-neuronal than neuronal H3K79me2 enrichment when compared to whole cortex, pointing the importance of non-neuronal for future studies (Figure 2J, FDR<0.05). Neurons specifically, showed a total of 0.35% (P28) and 7.1% (P51) of genes with differential H3K79me2 gene-body occupation (FDR<0.05, Figure 2D, Tables S3). Interestingly, H3K79me2 occupation increased from P28 to P51 in CTLs, but EE amplified this effect by affecting ∼2x more genes (Figure 2K, FDR<0.05, Table S3).

We revisited our previous findings in sorted neurons by addressing chromatin accessibility and gene-body epigenetic profiling (Figures 2C-D). Similar to whole cortex, we observed increased chromatin accessibility sites in enhancers and promoters after EE in NeuN^+^ cells (FC_neurons_=1.61X, Figure S4E). But differential sites represented around 0.01% and 0.006% respectively, a lower rate when compared to whole cortex (FDR<0.05, Figure 2B, C; Figure S3C, Table S2). As NeuN+ marker is specific for a broad number of different neurons (Figure 2I), we extended our study to sorted pyramidal neurons overexpressing a yellow fluorescent protein (YFP) under the control of the *Thy1* gene promoter (Method Details, Figure S3D). We found strongly increased chromatin accessibility upon EE (0.36% of enhancers, 0.39% of promoters), similar to the proportions in whole cortex (at FDR<0.05, Figures 2B-C; Figure S3E, F, Table S3).

Our findings confirm that EE leads to increased chromatin accessibility in whole cortex, in NeuN^+^ neurons, and more specifically in pyramidal neurons. We further confirm that these differential accessible enhancers are active forebrain enhancers at P0 and active in pyramidal neurons both in mouse and human (Figure S3G). Particular genes could be linked to human cognition in the context of schizophrenia and autism, such as *Nrg3* and *Ank2* respectively (Kao et al., 2010; Yang et al., 2019)(Figure S3H).

Because higher chromatin accessibility may allow increased TF binding, we ran a transcription factor binding site (TFBS) footprint analysis on whole cortex and NeuN^+^ populations using Centipede which screens for all putative TFBS (Pique-Regi et al., 2011). We confirmed that more TFs were significantly bound in EE compared to CTLs, such as Lhx3, AP1, Nr5a2 and Phox2B (Figure S3I; Table S2). Interestingly, we observed that CTCF displayed one of the strongest instances bound in EE-stimulated neurons, similar to CTCF ChIPseq results (p-adj<0.05, Figure 2C). This finding supports the idea that EE leads to increased genomic insulation and plays a role in genome organization during postnatal development.

Overall, we found that EE in neurons recapitulates cortical results inducing increased chromatin accessibility and CTCF binding. However in neurons, H3K79me2 increased upon EE which was not observed in the poised state of whole cortex. Noteworthy, GO terms of neuronal EE-induced changes show again genes associated with learning and memory targeting glutamatergic transmission predominantly, but also GABAergic and cholinergic transmission (Figure 2L, Figures S3J, K, L, Table S2).

### EE stimulation prompts 3D genome changes

The described EE-related changes implicate that the epigenome plays an important role in 3D genome organization (Guo et al., 2015; Rao et al., 2014). The evidence of increased chromatin accessibility and CTCF binding may suggest that environmental stimuli impact higher-order genome organization. To further assess chromatin interactions in sorted neurons (NeuN^+^), we performed Hi-C experiments to explore the 3D genome organization upon EE. By comparing *intra-*chromosomal interactions at 100 kilobase (kb) resolution, we determined significant changes: 94 interactions increased and 544 decreased upon EE stimulation (FDR<0.05, Figure 3A, Table S7). A decrease of *intra-*chromosomal interactions was also observed when calculating the number of chromatin loops using HICCUPs between CTL and EE (FDR<0.05, Figure 3B)(Durand et al., 2016a). We found differential *intra-*chromosomal bins to be clustered in particular chromosomal regions, specially involving chromosomes 8 and 14 (Figure 3C). In these regions, we find genes such as the synaptic-linked gene *Ngr1* linked to cognitive function improvement (Chen et al., 2008; Xu et al., 2016); and the synaptic vesicle exocytosis regulator *Cadps* (Sadakata et al., 2007).

**Figure 3.**
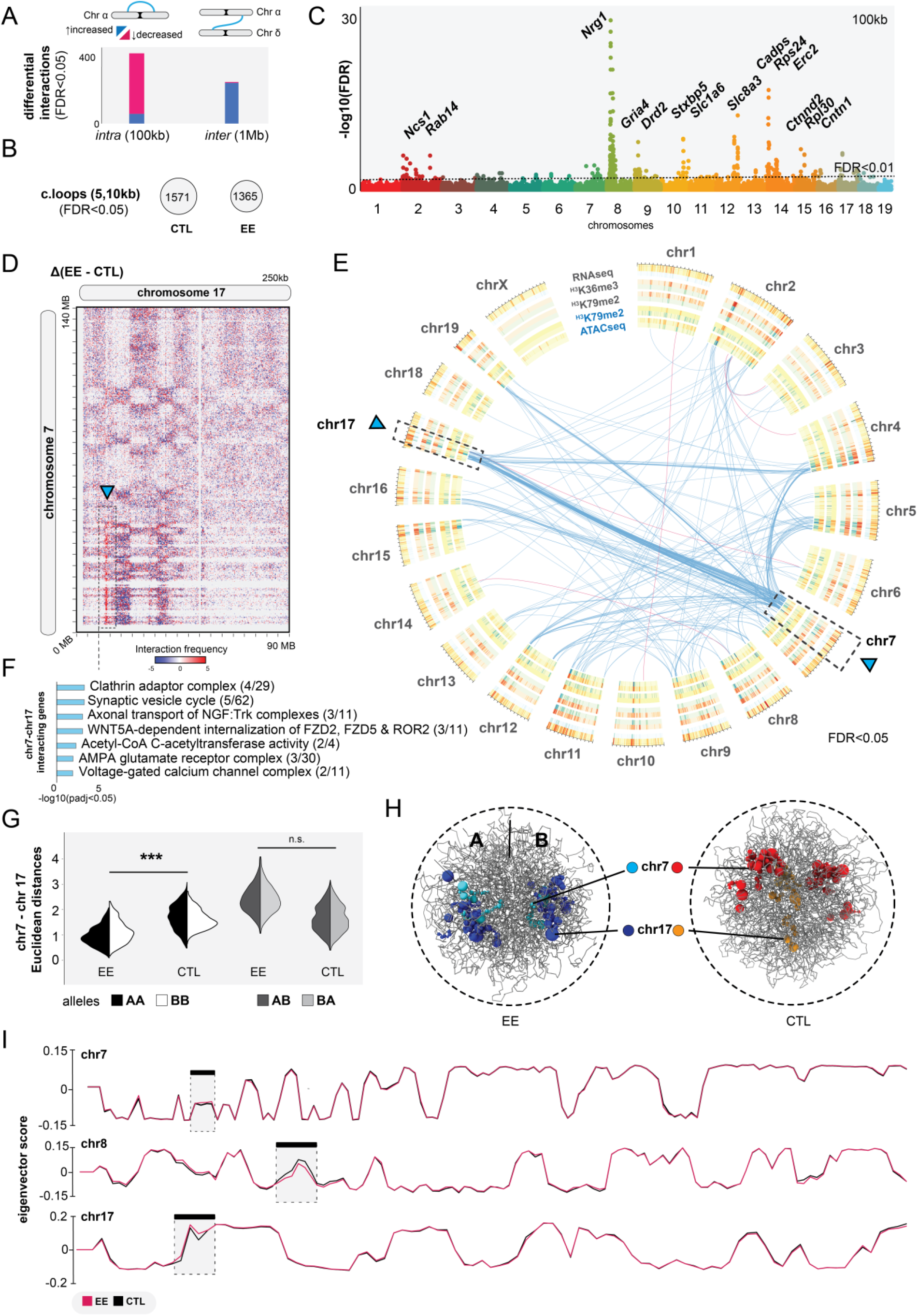
3D genome interaction changes upon EE. **A)** Differential analysis of *intra and inter*-chromosomal interactions at 100kb and 1Mb respectively, (FDR<0.05). **B)** Significant chromatin loops computed with HICCUPs at 5 and 10kb resolution (FDR<0.05). **C)** Manhattan plot of differential *intra*-chromosomal interactions at 100kb. **D)** Juicebox heatmaps at 250kb showing the extraction of EE versus CTL of *inter*-chromosomal interactions. **E)** Circos-plot of differential *inter*-chromosomal interactions (blue arcs-increased interactions, pink-decreased) together with concentric bedfiles representing the differential analysis of ATACseq, H3K79me2, H3K36me3 and RNAseq at 1MB using Diffreps (increased regions upon EE = blue, decreased = red). **F)** GO analysis of genes in the differential *inter-*chromosomal interactions at 1MB upon EE stimulation (p-adj <0.05 Bonferroni-step down). **G, H)** *In silico* chrom3D models for EE and CTL samples showing significant increase of inter-chromosomal interactions. **I)** A/B compartmentalization measured by eigenvector scores in chromosomes 7, 8 and 17.

While these *intra*-chromosomal contacts were in the focus of chromatin biology in recent years, contacts between different chromosomes (*inter-*chromosomal) also occur and are involved in important biological functions (Maass et al., 2019; Monahan et al., 2019a; Quinodoz et al., 2018), but they remain less studied. Therefore, we asked if EE is associated with large scale changes in genome organization by tracing chimeric *inter-*chromosomal Hi-C reads. Indeed, we identified 241 increased and 40 decreased interactions at 1 megabase [Mb] resolution (FDR<0.05, Figure 3A, Table S7). We determined a significant accumulation of *inter-*chromosomal contacts between chromosome 7 and 17 in EE *versus* CTL (Figure 3D-F, arrow). We observed that these differential interactions form a clear genome architectural stripe, also termed Greek islands (Monahan et al., 2019b)(Figure 3D). Among bins involving these two chromosomes, we found relevant genes associated with the synaptic vesicle cycle, such as *Ap2a*[1-2] being important for AMPAR endocytosis (DaSilva et al., 2016), *Acat[2-3]* acetyl-CoA C-acetyltransferase playing a key role in neuronal metabolism (Ronowska et al., 2018), and *Cacng8* modulating AMPAR receptor complexes in the plasmatic membrane which are important for synaptic plasticity (Maher et al., 2016) (Figure 3F, Table S7). We validated our differential analysis by an independent method, called Chrom3D, and generated *in silico* models for pooled EE and CTL samples detecting significant proximity of chromosomes 7 and 17 (Paulsen et al., 2017) (Figure 3G, H). We further corroborated the previous association of *inter-*chromosomal changes with gene-activity by studying chromatin compartmentalization (Lieberman-Aiden et al., 2009). As expected, A/B compartments do not change between CTL and EE, except for local changes in the strongest hubs of both *intra-* and *inter-*chromosomal contacts (chromosomes 7, 8, and 17), pointing to EE-related local chromatin changes in specific regions associated with active epigenetic modifications and gene expression changes (Figure 3I).

### EE causes coordinated regulatory changes that cluster in *inter*-chromosomal interactions implicated in memory-related functions

The multiple ‘omics’ datasets to study the molecular basis of EE allowed us to conduct an intersection of all EE-induced changes determined in this study (Figure 4A). Interestingly, GO analysis revealed synapse organization as the strongest ranked term (p-adj<0.05, Figure S4A). We decided to explore this finding further using SynGO synaptic gene curator tool to identify overrepresented genes (hits > 4 showing intersection in different sets, Figure 4B) (Koopmans et al., 2019). We found a significant enrichment of postsynaptic and presynaptic genes, particularly targeting the glutamatergic synapse (Figure 4C, D).

**Figure 4.**
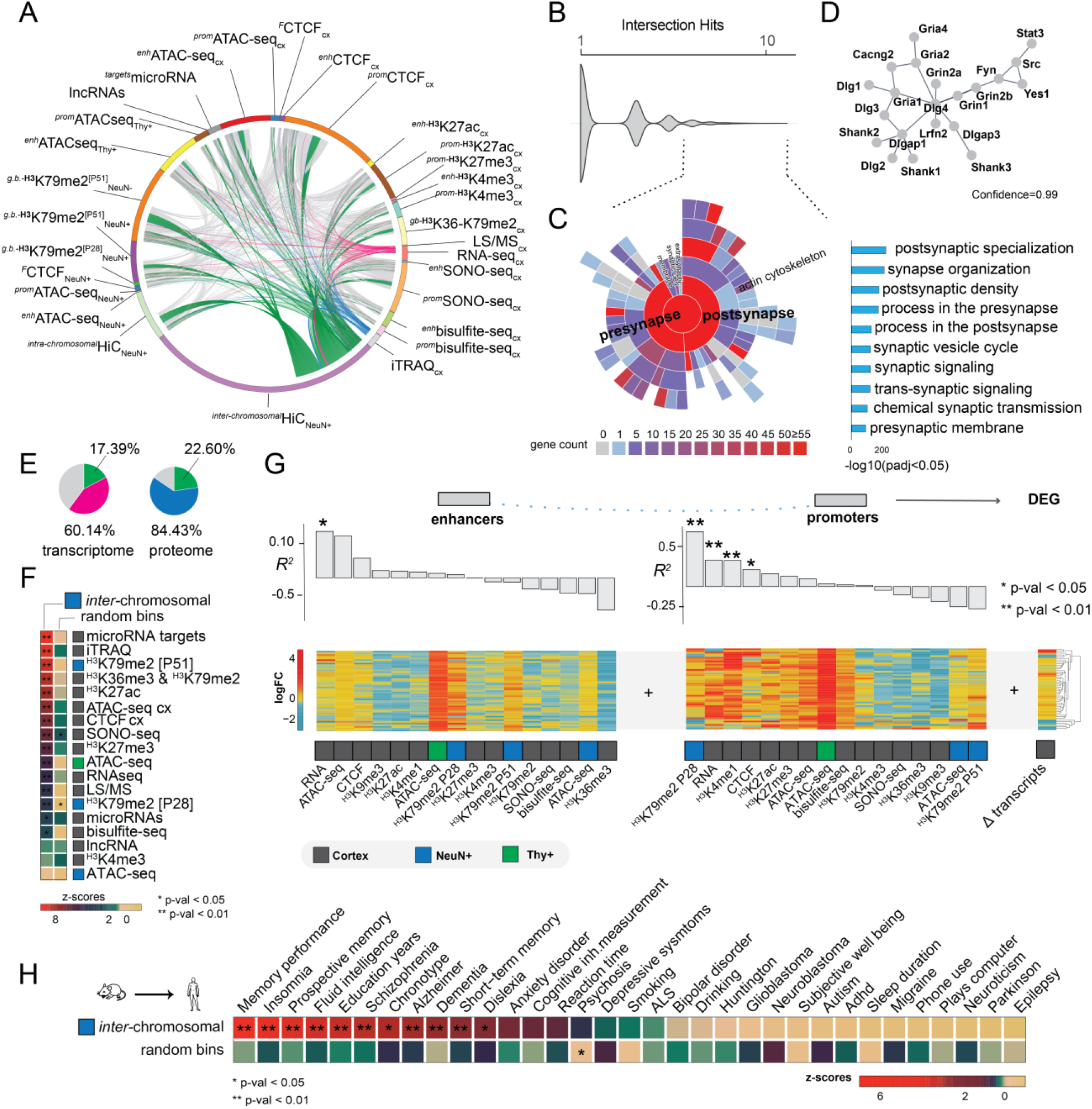
Data integration and EE implications in brain cognition. **A**) Full intersection of differential changes induced by EE (FDR<0.05). Pink arcs -differential expressed genes intersected with the rest of the data, blue proteomic, and green *inter*-chromosomal changes. **B**) Intersection hits plot, representing the number of times each gene is represented in the current study. Dashed lines -genes > 4 times intersected. **C**) SynGO analysis showing the enrichment of the most intersected genes which represent postsynaptic and presynaptic genes (right bar-plot, p-adj<0.05). **D)** String-db analysis interactome at 0.99 confidence of the most intersected genes. **E**) Transcriptomic and proteomic changes represented in other differential sets at FDR<0.05. “Pink + green” and “blue + green” -total percentages of transcriptomic and proteomic changes found in other differential datasets, where green specifically represents the portion of these changes found in inter-chromosomal changes. **F)** Differential *inter*-chromosomal changes association with the rest of the marks (N_permutations_= 100k, ** p-value<0.01, * p-value <0.05). **G**) Pearson translation efficiency of epigenetic marks in enhancers and promoters into differentially expressed genes (DEG). **H**) Differential *inter*-chromosomal changes association with human brain GWAS traits (N_permutations_= 100k, ** p-value<0.01, * p-value <0.05).

Further analysis of our merged data showed that about 60% of transcriptomic and 84% of proteomic changes are found in our other datasets, whilst 20% of changes were determined by EE-induced *inter*-chromosomal changes (Figure 4E). This enrichment together with the prior observation that differentially expressed genes tend to cluster in specific *inter*-chromosomal bins (Figure 3E), led us to the hypothesis that EE mainly induces changes locally in the genome, especially where active transcription occurs. To test this, we permuted the background genome at 1Mb resolution 100k times and calculated the likelihood of differential *inter*-chromosomal interactions to be associated with the differential epigenomic, transcriptomic, and proteomic changes found in the rest of the study. Strikingly, we observed that microRNA target genes, proteins (iTRAQ MS data), and gene-body associated histone marks were the strongest associated features within the specific *inter*-chromosomal hubs (Figure 4F). This finding confirms that EE orchestrates local changes of the nuclear architecture, especially inter-chromosomal communication.

Our intersection and permutation analysis indicated that chromatin conformation might connect the epigenome with the molecular phenotypes. We now asked how different marks influence others by estimating the linear dependency of EE-induced enhancers and promoters with transcriptomic and proteomic changes by Pearson and Spearman correlations (Figure 4G, Figure S4B, Method details). Transcription in enhancers ranked first and is the most correlated feature with transcriptomic and proteomic changes, whilst H3K79me2 correlated most at P28 in promoters (Figure 4G). These findings indicate that transcripts and proteins observed at P51 are dependent on earlier stages of postnatal neuronal development, thereby underlining the temporal aspect of the critical period.

The cognitive and behavioral effects of EE are similar in mice and human (Ball et al., 2019). Thus far, it is unclear if the molecular effects of EE that we found in mice can be retrieved in human. Therefore, we addressed the local changes in 3D genome organization in the human genome. We performed a lift-over of the differentially interacting *inter*-chromosomal bins at 1Mb from mouse to the human genome and ran a permutation analysis to test the association with 33 genome wide association studies (GWAS) relevant for human brain traits (Table S8). We observed that the top associated traits were memory-related, such as memory performance (p-val<0.01, Figure 4H). These findings not only validate our previous results, but point to conserved mechanisms between mouse and human that drive EE-related molecular effects by epigenomic, transcriptomic, and proteomic changes locally in specific regions of the genome that are important for both human and mouse cognition.

## Discussion

We have characterized the regulatory response to EE by using omics both in whole cortex tissue and in two neuronal cell populations and provide a valuable resource for other scientists. We demonstrate that EE induces coordinated changes of gene-regulatory networks that involve epigenetics and genome organization to adapt to constant cognitive stimulation and social interaction. EE induced increased enhancer and promoter chromatin accessibility in neurons, corroborating previous studies showing increased open chromatin upon invasive neuronal stimulation (Fernandez-Albert et al., 2019; Koberstein et al., 2018b; Su et al., 2017). These studies also highlighted the importance of CTCF shaping the 3D genome during postnatal development for memory formation (Kim et al., 2018; Sams et al., 2016). Here, we demonstrated that CTCF tends to bind preferentially to synaptic-associated genes upon EE, particularly glutamatergic associated pathways.

Our results also revealed, that gene body marks show differential activity in distal active enhancers upon EE, pointing to a potential role for these marks at transcriptionally active enhancers (Zentner et al., 2011). This is conform with the finding some DNA methyltransferases depend on H3K36me3 to exert their function at enhancers (Rinaldi et al., 2016). We also observed that active transcription in regulatory regions during early stages of postnatal neuronal development may influence local transcriptomic and proteomic changes at later stages. Furthermore, studying gene body marks in sorted neuronal populations allowed us to identify the molecular effects during the postnatal critical period, reflected by a constant increase in H3K79me2 occupation. We found it exacerbated in an experience-dependent manner, with a greater number of increased binding sites in EE compared to CTL samples across time.

Despite the caveat of cell heterogeneity potentially skewing observations in whole tissue-related experiments, particularly involving epigenetic gene body marks, it has been shown that these marks can be anticorrelated with expressed genes during aging (Pu et al., 2015). Furthermore, we have shown by cell-deconvolution the importance of other cell types which are often ignored in learning-memory studies. Additionally, EE-induced directional regulation particularly of epigenetic marks could reflect the discrepancy occurring upon cognitive stimulation, such as pruning and synaptic tuning, where both synaptic strength and loss are part of the learning process during postnatal development (Stephan et al., 2012; Turrigiano, 2008).

Furthermore, by applying Hi-C to neurons, we elucidated for the first time *intra-* and *inter*-chromosomal interactions sensitive to EE. Especially the mnemonic *inter-*chromosomal 3D conformation map with its major *inter-*chromosomal hub involving chromosomes 7 and 17 shows that the environmental stimulus EE affects local epigenomic regulation and chromatin interactions in a coordinated manner. EE-induced changes relate to both synapse strengthening and pruning genes, affecting cytoskeletal rearrangements and ECM associated genes (Smagin et al., 2018; Wright and Harding, 2009). These synaptic rearrangements need pathways such as Rho, GPCR, and PKC/Akt signaling which we found enriched in our study, with special enrichment of Wnt signaling (Hu et al., 2013; Lichti et al., 2014; Tan et al., 2017).

Our results indicate that environmental cues, particularly social interactions and constant EE stimulation, modulate epigenomic and 3D genome landscapes in a coordinated manner to achieve EE-cognitive improvement.

## Acknowledgments

We acknowledge support of the Spanish Ministry of Economy and Competitiveness, ‘Centro de Excelencia Severo Ochoa 2017-2021’, SEV-2016-0571, the CERCA Programme / Generalitat de Catalunya and Jerome Lejeune Foundation. Swiss National Science Foundation Fellowship (PBLAP3_136878) and Co-funded by Marie Curie Actions to CNH. This research was supported by the Social Sciences and Humanities Research Council of Canada (NFRFE-2018-01305). Resources for analyses conducted by S.E.G. were partially supported by U.S. National Institutes of Mental Health Funds R01MH104341 and R01MH117790. Special thanks to the “Centre for Genomic Regulation Core Facilities”: Irene González Navarrete, María Angustias Aguilar Morón, Anna Menoyo Vilalta, Núria Andreu Somavilla and Jochen Hecht from the Genomics Facility and Òscar Fornas, Eva Julià Arteaga, Erika Ramírez Bautista and Alexandre Bote Tronchoni from the CRG FAC-sorting Unit. Also, we would like to thank to Domenica Marchese for helping with microRNA validation and Jekaterina Kokatjuhha and Mattia Bosio for their computational help support.

## METHODS

- **KEY RESOURCES TABLE**
- **LEAD CONTACT AND MATERIALS AVAILABILITY**
- **EXPERIMENTAL MODEL AND SUBJECT DETAILS**
- **METHOD DETAILS**
  ▪ Nuclei acid extraction
  ▪ Nuclei isolation
  ▪ FAC-sorting
  ▪ Whole genome sequencing
  ▪ Chromatin accessibility
  ▪ Chromatin immunoprecipitation
  ▪ Transcriptomcs
  ▪ Proteomics
▪ Chromatin interactions: *in-situ* Hi-C
- **QUANTIFICATION AND STATISTICAL ANALYSIS**
- **DATA AND CODE AVAILABILITY**

## KEY RESOURCES TABLE

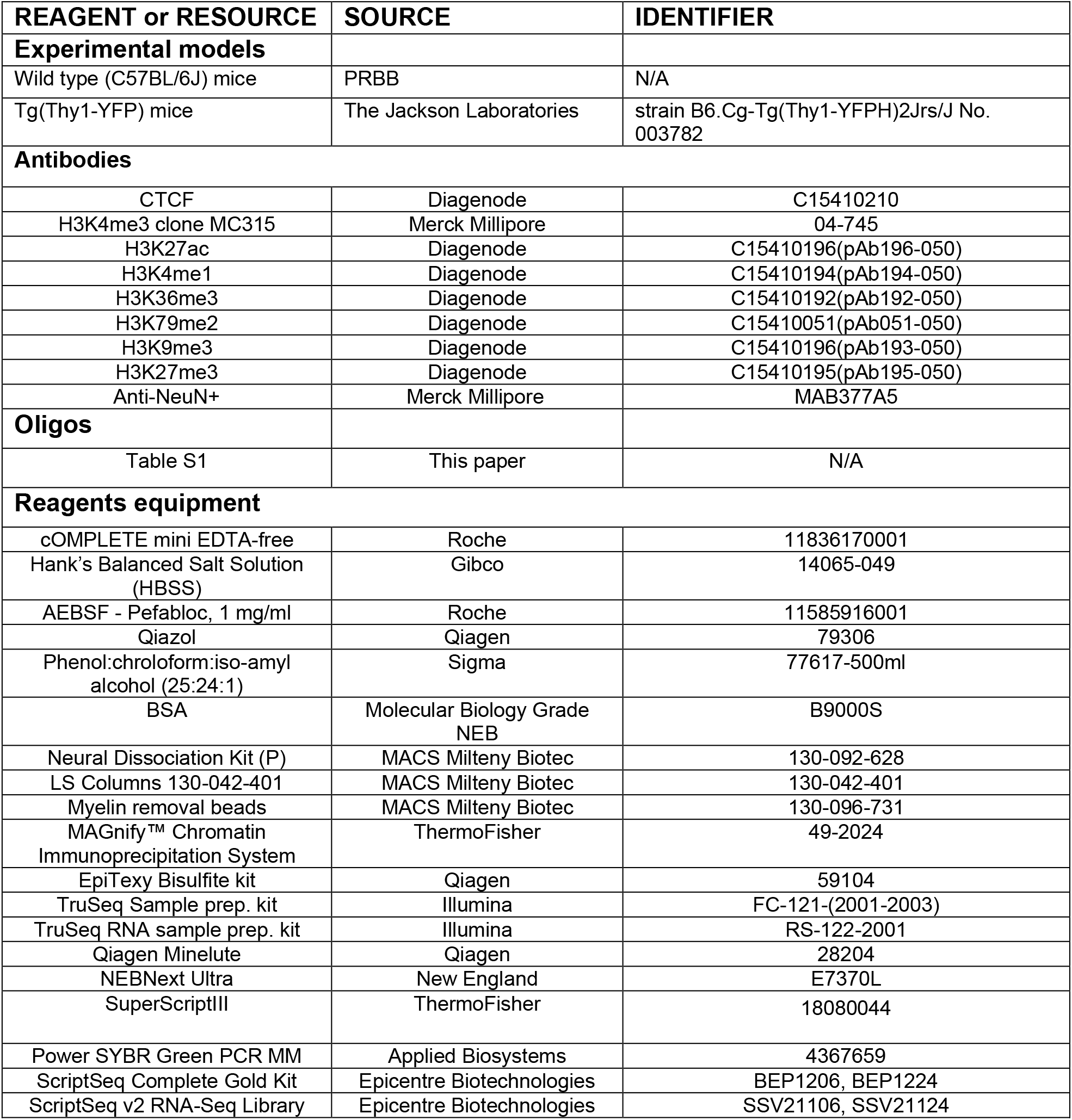
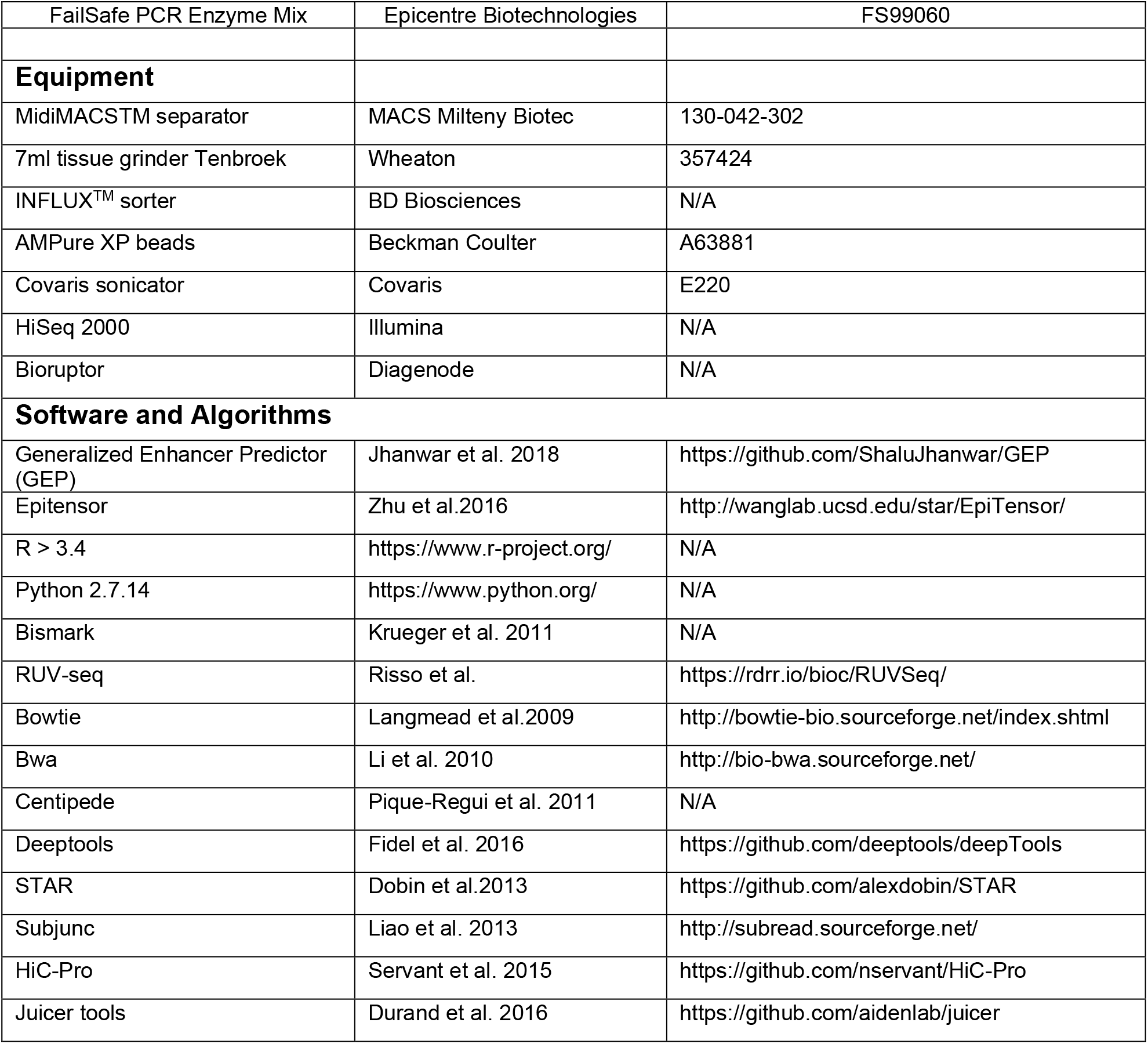

### LEAD CONTACT AND MATERIALS AVAILABILITY

Further information and requests for resources and reagents should be directed to the Lead Contact Sergio Espeso-Gil. This study did not generate any unique reagents.

### EXPERIMENTAL MODEL AND SUBJECT DETAILS

All experimental procedures were approved by the local ethical committee (Procedure Code: ISA-11-1358). Wild type mice (C57BL/6J) and Tg(Thy1-YFP) (strain B6.Cg-Tg(Thy1-YFPH)2Jrs/J No. 003782; The Jackson Laboratories) were kept and bred according to local (Catalan law 5/1995 and Decrees 214/97, 32/2007) and European regulations (EU directives 86/609 and 2001-486).

Upon weaning at three weeks of age, female pups were separated into control and enriched environment (EE) experimental groups. Cortical data analysis derived from the same EE protocol that was used by Pons-Espinal et al. 2013, where behavioral studies (Morris water maze testing) showed successful EE treatment. Briefly, control mice were kept under standard conditions with two individuals per cage, while EE consisted of keeping 8 mice in a large cage with toys, houses and tunnels, which were changed every three days for novelty. Both groups had unlimited access to food and water at all times. At five (P28) or eight weeks of age (P51), mice were euthanized by carbon dioxide, and the cerebral cortex was dissected within one minute of death. The tissue was immediately flash frozen in liquid nitrogen. In the case of Tg(Thy1-YFP) animals, the tissue was immersed in Hank’s Balanced Salt Solution (HBSS 1X, Gibco 14065-049) before proceeding with the sample preparation for the FAC-sorting (see section below). For each condition and replicate, we pooled the cortices of five mice with some exceptions: mice for *insitu-*HiC where single replicas as well as Thy-YFP mice for ATACseq (*N*_cortex_= 20 mice in 4 bioreplicates; *N*_sorted_= 30 mice in 6 bioreplicates, *N*_NeuN+_HiC_= 4 mice in 4 bioreplicates, *N*_Thy+_=4 mice in 4 bioreplicates). The frozen cortices for pooled animals were ground together in a frozen mortar containing liquid nitrogen, to obtain a fine powder of pooled cortex tissue. The powder was aliquoted and stored at -80°C until further use.

### METHOD DETAILS

#### Nucleic acid extraction

DNA was extracted using Phenol:chroloform:iso-amyl alcohol (25:24:1) according to manufacturer guidelines (Sigma 77617-500ml). RNA was extracted using Qiazol total RNA (Qiagen Cat No:79306) kit according to the manufacturer’s instructions. The RNA was quantified by Qubit ® 2.0 Fluorometer (Life Technologies) and the quality was assessed using a Nanodrop 2000c (Thermo Scientific) and a 2100 Bioanalyzer (Agilent Technologies, CA, USA).

#### Nucleus isolation

To obtain fresh nuclei, ground frozen tissue was resuspended in tissue lysis buffer (1x PBS containing 0.1% Triton X-100, 1x Complete Protease inhibitor cocktail tablette (cOMPLETE mini EDTA-free, Roche Cat No.11836170001) and 1 mg/ml AEBSF (Pefabloc, Roche Cat No.11585916001)) and dissociated by 60-90 strokes in a glass douncer (7ml tissue grinder Tenbroek, Wheaton Cat No. 357424). Nuclei were counted using a hemacytometer and constantly checked under a microscope (Leica DM-IL).

#### FAC-sorting

Two different procedure were performed: 1) <u>Sorting total neurons using NeuN (Rbfox3) marker</u> and 2) Sorting pyramidal neurons in Tg(Thy1-YFP) mice. Briefly, after the nucleus preparation, nuclei were resuspended in 1ml of PBS-PI 1X (1X-PBS, 1x Complete Protease inhibitor cocktail (cOMPLETE mini EDTA-free, Roche Cat No.11836170001), 1 mg/ml AEBSF Pefabloc (Pefabloc, Roche Cat No.11585916001) and 0,1 mg/ml of BSA (BSA, Molecular Biology Grade NEB, Cat No.B9000S)). 1,5 μl of anti-NeuN, clone A60, Alexa Fluor ® 555 Conjugate (Merk Millipore Cat No. MAB377A5) was added to the solution and incubated at 4°C for 1.5h protected from light. The sample was centrifuged for 10 mins, 500g at 4°C and washed with 1ml of PBS-PI 1X. Next, 1μl of 4′,6-Diamidine-2′-phenylindole dihydrochloride was added (DAPI, Roche Cat. No. 10236276001) and the sample was given immediately to the FACS-sorting Facility. Samples were sorted at 12PSI in cold condition in an INFLUX sorter (BD Biosciences INFLUX™). The sorted samples were centrifuged for 40min at 700g at 4°C to collect the nuclei before proceeding with the desired technique. For sorting pyramidal neurons Tg(Thy1-YFP), animals were dissected and tissue was immediately submerged in Hank’s Balanced Salt Solution (HBSS 1X, Gibco 14065-049). Brain samples were dissociated using the Neural Dissociation Kit (P) (MACS Milteny Biotec Cat.No. 130-092-628; LS Columns Cat.No 130-042-401; Myelin removal beads Cat No. 130-096-731; MidiMACSTM separator Cat.No. 130-042-302), according to manufacturers’ instructions. Cells were sorted using an INFLUX™ sorter (BD Biosciences INFLUX™). After sorting, samples were centrifuged for 40mins at 700g at 4C to collect the nuclei before proceeding with the desired technique.

#### Whole genome bisulfite-sequencing

WGBS was performed by CNAG Genome Facility on two independent sets of biological replicates. Briefly, genomic DNA (1–2 μg) was spiked with unmethylated λ DNA (5 ng of λ DNA per microgram of genomic DNA; Promega). DNA was sheared by sonication to 50–500 bp in size using a Covaris E220 sonicator, and fragments of 150–300 bp were selected using AMPure XP beads (Agencourt Bioscience). Genomic DNA libraries were constructed using the Illumina TruSeq Sample Preparation kit following Illumina’s standard protocol. DNA was treated with sodium bisulfite after adaptor ligation, using the EpiTexy Bisulfite kit (Qiagen), following the manufacturer’s instructions for formalin-fixed, paraffin-embedded tissue samples. Two rounds of bisulfite conversion were performed to ensure a conversion rate of >99%. Enrichment for adaptor-ligated DNA was carried out through seven PCR cycles using PfuTurboCx Hot-Start DNA polymerase (Stratagene). Library quality was monitored using the Agilent 2100 Bioanalyzer, and the concentration was determined by quantitative PCR with the library quantification kit from Kapa Biosystems. Paired-end DNA sequencing (2 × 100 bp) was then performed using the Illumina HiSeq 2000 platform.

#### Chromatin accessibility

Open chromatin studies were performed by ATAC-seq and SONO-seq procedures. Briefly, ATAC-seq was performed with minor modifications from Buenrostro et al.(Buenrostro et al., 2013). 100’000 nuclei were treated with 2.5 µl Tn5 at 37°C for 30 minutes, followed by cleanup on a Qiagen Minelute column. Fragments >1kb in size were removed using AmpureXP beads (Agencourt AMPure XP, Beckman Coulter Cat.No. A63881). DNA fragments were amplified by 11 cycles of PCR with custom adapter primers from Buenostro et al. (Buenrostro et al., 2013). PCR reactions were cleaned up with AmpureXP beads, quantified by Qubit and quality controlled by Bioanalyzer (Agilent Technologies). SONO-seq consists of isolating and fragmenting crosslinked chromatin, before reversing crosslinks, purifying the DNA and preparing it for sequencing(Auerbach et al., 2009). Chromatin was fragmented by sonication using the same COVARIS specifications as for ChIP-seq (see above) to obtain a median fragment size of 200 bp. Sequencing libraries were prepared using the NEBNext Ultra (New England Cat.No. E7370L) kit according to the manufacturer’s protocol. Both techniques were sequenced on a HiSeq2000 sequencer (Illumina).

#### Chromatin Immunoprecipitation: ChIP-seq

Nuclei obtained in section 1.2. were cross-linked with 0,5% formaldehyde (Sigma F8775-25ml) for 5 minutes at room temperature (RT). Residual formaldehyde was quenched by addition of glycine (MAGnify™ Glycine P/N 100006373) to a final concentration of 0.125M and incubation for 5 minutes at RT. Nuclei were pelleted by centrifugation at 500g during 10 min at 4°C and resuspended in 300ul lysis buffer (MAGnify™ Chromatin Immunoprecipitation System, Cat no.49-2024). Chromatin was fragmented by sonication in a Covaris S2 (Duty Cycle: 20, Intensity: 8, Cycles per Burst:200, for 15mins (histone marks), for 25 mins (FACS-sorted nuclei)) to a median size of 200 bp, aliquoted and stored at -80°C until further use. For non-histonic proteins such CTCF, no nuclei preparation was performed. Homogenized tissue was crosslinked with 0,5% formaldehyde for 10 minutes at RT, quenched and fragmented as above (Duty Cycle: 5, Intensity: 2, Cycles per Burst: 200, Time: 25mins). Chromatin immunoprecipitation was performed using antibodies against histone modifications (H3K27ac, H3K4me3, H3K4me1, H3K79me2, H3K36me3, H3K27me3, H3K9me2 and CTCF) with the MAGnify™ Chromatin Immunoprecipitation System, (Invitrogen Cat no.49-2024), according to manufacturer’s instructions. For whole cortex and NeuN histonic ChIP-seq a total amount of 50k nuclei was used per ChIP (∼330ng), 700k nuclei (∼4μg) for non-histonic ChIP-seq. Recovered ChIP DNA was used to construct sequencing libraries, using the NEBNext Ultra (New England Cat.No. E7370L) kit according to the manufacturer’s protocol, and sequenced on a HiSeq2000 sequencer (Illumina). The quality of the ChIP-seq was determined by qPCR, using positive and negative primers to detect the regions where the histones should be placed in the genome (Table S1). Primers were diluted to a final concentration of 300ng in Power SYBR Green PCR MM (Applied Biosystems Cat.No 4367659). Samples were run in a Applied Biosystem qPCR system (7900 HT Fast Real-Time PCR System) as follows: 50°C/2min, 95°C/10min, 40 cycles of 95°C/15s, 60°C/1min, 95°C/15s, 60C-15s and 95-15s.

#### Transcriptomics

Transcriptome study involved both poly-A RNA, directional RNA and small RNA libraries. Poly-A RNA sequencing libraries were prepared from total RNA using the TruSeq™ RNA sample preparation kit (Illumina Inc., Cat.No. RS-122-2001) according to the manufacturer’s protocol. Directional RNA libraries were prepared using the ScriptSeqTM Complete Gold Kit (Human/Mouse/Rat) (Epicentre Biotechnologies), according to the manufacturer’s protocol. Briefly, 3 μg of total RNA were depleted of both cytoplasmic and mitochondrial rRNAs using the Ribo-Zero™ Gold rRNA Removal Reagents. The total rRNA depletion of the samples was confirmed on a 2100 Bioanalyzer RNA 6000 Pico Chip. For the library preparation we used 50 ng of Ribo-Zero-treated RNA as starting material for the ScriptSeqTM v2 RNA-Seq Library Preparation Kit, followed by amplification by 12 cycles of PCR, using the FailSafeTM PCR Enzyme Mix (Epicentre Biotechnologies) before purification with AMPure XP beads (Agencourt, Beckman Coulter Cat.No. A63881). Both the directional mRNA and the Poly-A RNA libraries were sequenced in paired end mode with read length 2×101bp on a HiSeq2000 (Illumina, Inc) following the manufacturer’s protocol. After computational analysis, we validated 21 differential expressed genes in a new batch of biological replicates following the method of Schmittgen & Livak(see Table S4)(Schmittgen and Livak, 2008). To validate the data generated in the RNA-seq analysis, 1 ug of the sequenced RNAs were used to prepare cDNA with SuperScriptIII (Invitrogen) according to manufacturer’s instructions, cDNAs were normalized to 100ng/ul. RT-PCRs for the alternative splicing events were performed using oligos annealing to the adjacent constitutive exons and performed under standard conditions; 2% agarose gels were used to resolve the different bands. Image J software was used for quantification of the observed bands and determination of the PSIs for each event. Small RNA libraries were generated using the TruSeq (Illumina, Cat.No. RS-122-2001) kit according to the manufacturer’s protocol. The resulting 22bp insert libraries were sequenced on a HiSeq2000 sequencer (Illumina), yielding 15-20 million reads per sample. After the analysis, we validated 10 miRNA following Chen et al. protocol with minor modifications: instead of using Taqman probes we designed our own RT-miRNA oligonucleotides and performed qPCRs with Power SYBR Green PCR MM (Applied Biosystems Cat.No 4367659, see Table S5)(Chen et al., 2005).

#### Proteomics

Samples were minced with RIPA-M buffer (1% NP40, 1% Sodium deoxycholate, 0.15M NaCl, 0.001M EDTA, 0.05 TrisHCl pH=7.5, 1X cOMPLETE Mini EDTA free, 0.01M NaF, 0.01M Sodium pyrophosphate, 0.005M β-glycerolphophate), sonicated with a Diagenode Bioruptor (cycles of 0.5min ON 0.5 min OFF, medium intensity during 5min). Samples were centrifuged during 10min 16000rmp at 4°C and precipitated with acetone at -20°C for 1 hour. Samples were pelleted by centrifugation during 10mins 16000rpm at 4°C, dried and resuspended in Urea/200mM ABC, sonicated again during 10min (cycles of 0.5min ON 0.5min OFF, medium intensity) and quantified prior to mass spectrophotometry injection following isobaric tags for relative and absolute quantitation (iTRAQ) or Liquid Chromatography/Mass-Spectophotometry (LC/MS).

#### *in situ*-HiC

Cerebral cortex samples from individual C57BL/6J mice (2 bio-replicates per EE and CTL conditions) were sorted using NeuN^+^ (Rbfox3+) as described above. After sorting, approximately 1 million of nuclei were used for in situ HiC following previous specifications(Rao et al., 2014). Libraries were sequenced on a HiSeq2000 (PE x 125bp) yielding approximately 300M of reads per sample.

### QUANTIFICATION AND STATISTICAL ANALYSIS

#### *In silico* identification of active enhancers

Active enhancers for EE and CTL cortex were predicted and annotated using a machine learning approach called Generalized Enhancer Predictor (GEP, https://github.com/ShaluJhanwar/GEP)(Jhanwar et al., 2018). The method performed classification of epigenetic patterns coming from cortical ATAC-seq, H3K4me1, H3K4me3, H3K27ac, H3K36me3 and H3K27me3 using Random Forest (RF) and Support Vector Machine (SVM) classifiers to build predictive models for identification of active enhancers. The total amount of enhancers used in the present study corresponded to the consensus of GEP prediction for EE, control and ENCODE data, resulting in 347112 enhancers (Table S1).

#### *In silico* prediction of chromatin interactions

Chromatin interactions were predicted from chromatin modifications using Epitensor with minor modifications to adapt the script to the mouse genome (Zhu et al., 2016). We used epigenetic data from following tissues from the mouse ENCODE project (forebrain, heart, hindbrain, kidney, liver, lung, midbrain, stomach)(Yue et al., 2014). As well as in-house generated data from cortex of animals with and without environmental enrichment. We used the following epigenetic marks: H3K9me3, H3K4me3, H3K4me1, H3K36me3, H3K27me3, H3K27ac, CTCF as well as RNA-seq coverage data. Chromatin accessibility was measured by DNase-seq for ENCODE tissues and ATAC-seq for in-house cortex samples. Promoter regions were defined using the *TxDb*.*Mmusculus*.*UCSC*.*mm10*.*knownGene* package in BioConductor as 1500 up- and 500 down-stream of any possible TSS. Active enhancers in each of the input tissues were defined by training a machine learning classifier on a list of validated enhancers using core epigenetic mark intensities as features (DNase, H3K27ac, H3K27me3, H3K36me3, H4K4me1, H3K4me3 as well as intensity ratio between the last two marks). The trained classifier was then applied to the epigenetic data from ENCODE tissues and local cortex datasets. To limit computation load, possible interactions were limited to intra-TAD interactions, which were based on the set of mouse cortex TADs (http://chromosome.sdsc.edu/mouse/hi-c/download.html)(Dixon et al., 2012). Differential activity in enhancers was annotated using *annotate*.*enhancers*.*with*.*genes*.*sh* utility (https://github.com/ophiothrix/enhancer.annotator)

#### Whole genome bisulfite-sequencing

Methylated CpGs were called from the raw reads using Bismark(Krueger and Andrews, 2011) following the user guide. Differential methylation analysis was carried out using bsseq (Hansen et al., 2012). Briefly, CpG methylation values were locally smoothed using the BSmooth function and CpGs that had a coverage of less than 5 reads in any of the samples were removed. We calculated the t statistic for the smoothed CpG values using the BSmooth.tstat function with paired design in local correction mode and the dmrFinder function was used to identify differentially methylated regions (DMRs) that contained multiple CpGs with a t statistic below -4.5 or above 4.5. DMRs located inside or within 5 kb of a gene or enhancer were annotated accordingly. Additionally, DMRs that overlapped enhancers, were annotated with the enhancer’s target gene according to the Epitensor predictions.

#### Chromatin accessibility

ATAC-seq libraries were aligned to mm10 using bwa-mem with predefined parameters(Li and Durbin, 2010). Duplicate read pairs were marked using the *MarkDuplicates* command from the Picard software suite (http://broadinstitute.github.io/picard). We used a peak-independent approach on the one hand using our predicted enhancers and promoters as defined above, and on the other hand a peak-dependent method using F-seq or MACS2 with default parameters with and without duplicates (Boyle et al., 2008; Zhang et al., 2008). We also used MACS2 with the shifted strategy with the following parameters: “--nomodel --shift -75 --extsize 150 --broad --keep-dup all”. Together with enhancer and promoter regions, each annotation was loaded into DiffBind(Ross-Innes et al., 2012) providing as background input SONO-seq chromatin (Ross-Innes et al., 2012). For SONO-seq libraries, the pipeline followed the same steps as the ChIP-seq (see bellow).

Footprinting analysis was done using the Centipede software using the core transcription factor binding motifs from the Jaspar database, (version 2016-03-02)(Khan et al., 2018; Pique-Regi et al., 2011). Instances of each motif in the genome were kept if the PWM score was superior or equal to 13. We created and used a mm10 mappability file to filter out instances that are located in regions of the genome with low mappability (gem-mappability-retriever (Marco-Sola et al., 2012)). A motif was considered bound if the Pvalue-Zscore-combined statistics was inferior to 0.05. A motif was considered differentially bound if the ANOVA p-value was inferior to 0.05. Individual instances of a motif were considered bound when their posterior probabilities were superior to 0.99.

#### Chromatin Immunoprecipitation: ChIP-seq

Samples were mapped to mouse mm10 (peak-independent) using Bowtie (Langmead et al., 2009) with “--quiet --sam --best --strata -m 1” parameters. Sam files were converted to bam files allowing only reads with mapping quality greater than 30 (“-q 30”). Files were visualized using bamCoverage utility from deepTools(Ramírez et al., 2016). We initially used a peak-dependent strategy as a benchmarking method and also as a tool to define our peak-independent approach (data not shown). The quality of ChIP-seq was assessed by the irreproducible discovery rate (IDR)(Li et al., 2011). The peak-independent method used in the present study consisted in the calculation of the coverage at defined regions as follows. The broad marks H3K36me3 and H3K79me2 were measured along the full gene body, defined as the distance from the TSS to the TES. Enhancer regions were provided by the GEP enhancer predictor in combination with ENCODE enhancers (Jhanwar et al., 2018)(Table S1). Promoter regions were defined as the interval from 1500bp upstream to 500bp downstream of the TSS. Reads were quantified by featurecounts (Liao et al., 2014) with the following parameters: “--ignoreDup --minReadOverlap 25 -Q 1 -O”. Counts tables were supplied to the batch effect corrector RUVseq for further differential analysis using edgeR (Risso et al., 2014; Robinson et al., 2010). A binned approach was used to determine differential changes in 1Mb bins to be associated with inter-chromosomal interactions hubs. This differential analysis was performed with Diffreps (Shen et al., 2013). Using the following parameters: “-pval 0.001 -frag 150 -window 1000”. Batch effects are a major issue in sequencing studies. In this study, we acknowledge the lack of bio-replicates in the cortical tissue by having a wide set of different techniques as well as a pooling strategy of animals per bio-replicate (5 individuals per sample). To minimize batch effects, we randomized the samples, applied standardized procedures and parallelized the experiments as much as possible. However, FAC-sorting experiments could not always be parallelized for different reasons (i.e. availability of the sorter). To remove batch effects, we decided to use the strategy of Russo et al. that was also used in data different from RNAseq (Koberstein et al., 2018b; Risso et al., 2014). A principal component analysis was the main criterion to evaluate each dataset and the requirement for batch correction. If a PCA on uncorrected data could clearly separate the conditions, we ruled that there was no need for batch correction. However, if required, we then selected the method(s) (among RUVg, RUVs, and/or RUVr) that were able to separate conditions and intersected the results with a FDR threshold of 0.1 to obtain a conservative final gene list of changes induced by EE. Coverage plots were produced using the function bamCoverage of deepTools to produce bigwig files (Ramírez et al., 2016) (v2.0 with Python 2.7.14). These files were supplied to the function computeMatrix using the “scale-regions” parameter that allows to plot the coverage profile using the plotProfile function along regions of interest. The cell-specificity of sorted populations was assessed by the tool MakerGeneProfile that assesses cell specificity using a curated single-cell mouse brain RNAseq database (Mancarci et al., 2017). We transformed gene-body assigned reads of H3K79me2 NeuN^+^ and NeuN^-^ into RPKM values and ran MakerGeneProfile to compare neuronal and non-neuronal enrichments to oligodendrocytes, astrocytes and pyramidal neurons specific markers. Gene ontology term enrichment analysis was performed using the Cytoscape tools clueGO and cluepedia, Metascape and SynGO (Bindea et al., 2009; Koopmans et al., 2019; Zhou et al., 2019). We performed the analysis individually for each histone mark or together, supplying upregulated and downregulated DBS separately. We used Bonferroni step down or Benjamini-Hochberg with p-value thresholds of 0.05 or 0.01 depending of the amount of data provided. In general, larger datasets needed more stringency in the statistical correction (Benjamini and Hochberg, 1995).

#### Transcriptomics

mRNA reads were aligned to mm10 using STAR with standard parameters (Dobin et al., 2013). Reads were counted using featureCounts by “gene_name” (Liao et al., 2014)”, batch corrected by RUV-seq before differential analysis in edgeR (Risso et al., 2014; Robinson et al., 2010). We also provide a splicing analysis results compiled in Table S4. To identify and quantify alternative splicing, we used *vast-tools v1*.*0*.*0* (Tapial et al., 2017). To increase effective read coverage at splice junctions, we pooled all replicates for polyA, respectively ribominus, samples using *vast-tools merge*. Differential splicing analysis was performed using *vast-tools compare*, comparing EE with Ctl samples in a paired manner, using the parameters --min_dPSI 10 --min_range 5 --p_IR --noVLOW. This resulted in respectively 40 and 53, cassette exons being up- or downregulated with EE (including 2 and 1 microexons (Irimia et al., 2014)]), 49 and 43 retained introns, 27 and 22 alternative 3’ss choices and 28 and 23 alternative 5’ss splice choices.

Regarding the lncRNA analysis, total RNA reads were aligned to mm10 using Subjunc splice-aware aligner with default settings (Liao et al., 2013), and reads overlapping exons were summarized at the gene-level to the corresponding genes using featureCounts (Liao et al., 2014). Read assignment to exons was carried out in a strand-aware manner, only fragments with both mates correctly aligned were considered and genomic regions with multiple overlapping exons on the same strand were disregarded. The count matrix was further filtered to retain only GENCODE long non-coding RNA (GRCm38.p5_M15). Reads with an average log2 CPM <0.9 across all samples were filtered out yielding 4472 genes. Normalization factors were calculated using the TMM method from edgeR package (Robinson et al., 2010). Observational-level weights were calculated using voom (Law et al., 2014) and used to fit gene-wise linear models (Smyth, 2004). Multiple testing p-value adjustment was performed using the Benjamini-Hochberg method (Benjamini and Hochberg, 1995)

Small RNA reads with homo-polymer and low PHRED scores were removed using FASTQ-Toolkit and a custom script. SeqBuster was used to remove adapters and align using miraligner.jar with mouse miRbase v18 annotation(Pantano et al., 2010). The small RNA dataset presented a strong batch effect which could not be corrected by RUVseq(Risso et al., 2014). We therefore devised an alternative strategy which consisted of filtering microRNAs for low read coverage (<50 counts), normalizing the libraries by read counts per million (rpm), before contrasting EE against CTL batch-wise and considering exclusively reproducible direction of change. Additionally, we calculated z-scores validating partially previous strategy (Table S5).

#### Proteomics

The analysis of iTRAQ data consisted in sorting the discrepancy (*δ* =φ/β) between both biological contrasts (φ = EE1/CTL1 and β = EE2/CTL2), following the premise that consistent results will be indicated by:

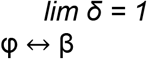

Discrepant results then will be considered as values far from *δ* =1. Based on this consideration we select all the proteins that follow the condition and we established a threshold of ±0.1. The protein expression values from the LC/MS were log2-transformed and loess normalized using the *normalize*.*ExpressionSet*.*loess* function from the BioConductor package AffyPML. Differential expression analysis was conducted with a standard limma (BioConductor) pipeline by calculating sample weights, fitting a linear model for each gene across all samples and calculating moderated F-statistics. Unadjusted p-values were used to rank the proteins (Smyth, 2004).

#### *In situ* Hi-C

For quality check, sequencing reads were mapped to the mouse reference genome assembly (mm10), artifacts were filtered, and library was ICED normalized using the Hi-C-Pro (Servant et al., 2015) (v2.9.0) (Table S7). For visualization we used the *hicpro2juicebox*.*sh* utility to convert the previous into a *.hic format to visualize heatmap interaction matrices in Juicebox(Durand et al., 2016b). Juicer_tools were used to calculate A/B compartments using eigenvector utility (Durand et al., 2016a). We called chromatin loops with HICCUP at 5 and 10kb of resolution (Durand et al., 2016b). For differential analysis, resulting interactions from Hi-C-Pro were splitted into intra and interchromsomal interactions, sex chromosomes and self-interacting bins were removed. Then piped into the edgeR wrapper RUVseq were RUVr for *intra* and RUVg for *inter* were used to normalized batch effects.

Required EE and CTL gtrack file to run chrom3D (Paulsen et al., 2017) was produced using the chrom3D wrapper automat_chrom3D utility (https://github.com/sespesogil/automat_chrom3D). Domains were called using Arrowhead (Juicer tools 1.7.6 (Durand et al., 2016a)). “--ignore_sparsity” parameter was used, and calls could be only produced at not lower than 10 kb. For the present study, we finally selected 5M iterations including the parameter “--nucleus” to force the beads to remain confined inside the designed radius: “-r 3.0”. Domain coloring was produced by automat_color (https://github.com/sespesogil/automat_chrom3D_colors) that allows to color any region of interest in the model. Both *in silico* models are available Table S7.

#### Data integration

We used Metascape to intersect the totality of the data. Some of the results were clumped as the maximum amount of sets allowed is 30(Zhou et al., 2019). Transcriptomic and proteomic changes percentages explain by the rest of the data were calculated out of the evidences table reported as Metascape result result (Table S8).

Linear dependency test was performed by using Pearson and Spearman correlations to test how enhancer and promoter activity of different marks influence differential changes observed in the transcriptome and proteome. Briefly, normalized counts by RPKM mapping into enhancers and promoters were first averaged by target gene name they interact with using EpiTensor (Zhu et al., 2016). Then a fold change was calculated of EE vs CTL samples. These values then are correlated with the corresponding fold changes found in differential RNA-seq and proteomic analysis. We used Spearman for proteomic changes as the scale of the fold changes were considerably different coming directly from iTRAQ data and the method is less sensitive for these outliers.

In order to ttest he the association of differential regions induced by EE with *inter*-chromosomal interactions we run a permutation analysis using the package *regioneR*. Both permutations shown in the study were performed with a total number of 100000 iterations. GWAS brain trait studies are collected in detail in Table S8.

### DATA AND CODE AVAILABILITY

Datasets are currently accessible upon request in the SRA repository: https://www.ncbi.nlm.nih.gov/sra/SRP154319.

Code present in this study can be found in the following repository: https://github.com/sespesogil/Environmental_enrichment

Browser data visualization is available here: http://epigenomegateway.wustl.edu/legacy/?genome=mm10&session=vs5yrq9qOX&statusId=1272141941

## SUPPLEMENTARY FIGURES

**Figure S1. Chromatin accessibility and epigenetic changes induce by EE in cortex homogenate**

**Figure S2. Overlap of differential binding sites due EE**.

**Figure S3. Transcriptional and translational changes due EE**.

**Figure S4. Chromatin accessibility and transcription-associated changes due EE in sorted populations**

### SUPPLEMENTARY TABLES

**Table S1. Annotations used in the present study**.

**Table S2. EE-induced chromatin accessibility**.

**Table S3. EE-induced epigenetic modifications**.

**Table S4. EE-induced coding transcriptomic changes**.

**Table S5. EE-induced non-coding transcriptomic changes**.

**Table S6. EE-induced proteomic changes**.

**Table S7. EE-induced 3D genome changes**.

**Table S8. Data integration**.

