## Supplementary Figures for "Environmental enrichment induces epigenomic and genome organization changes relevant for cognitive function"

### Figure S1

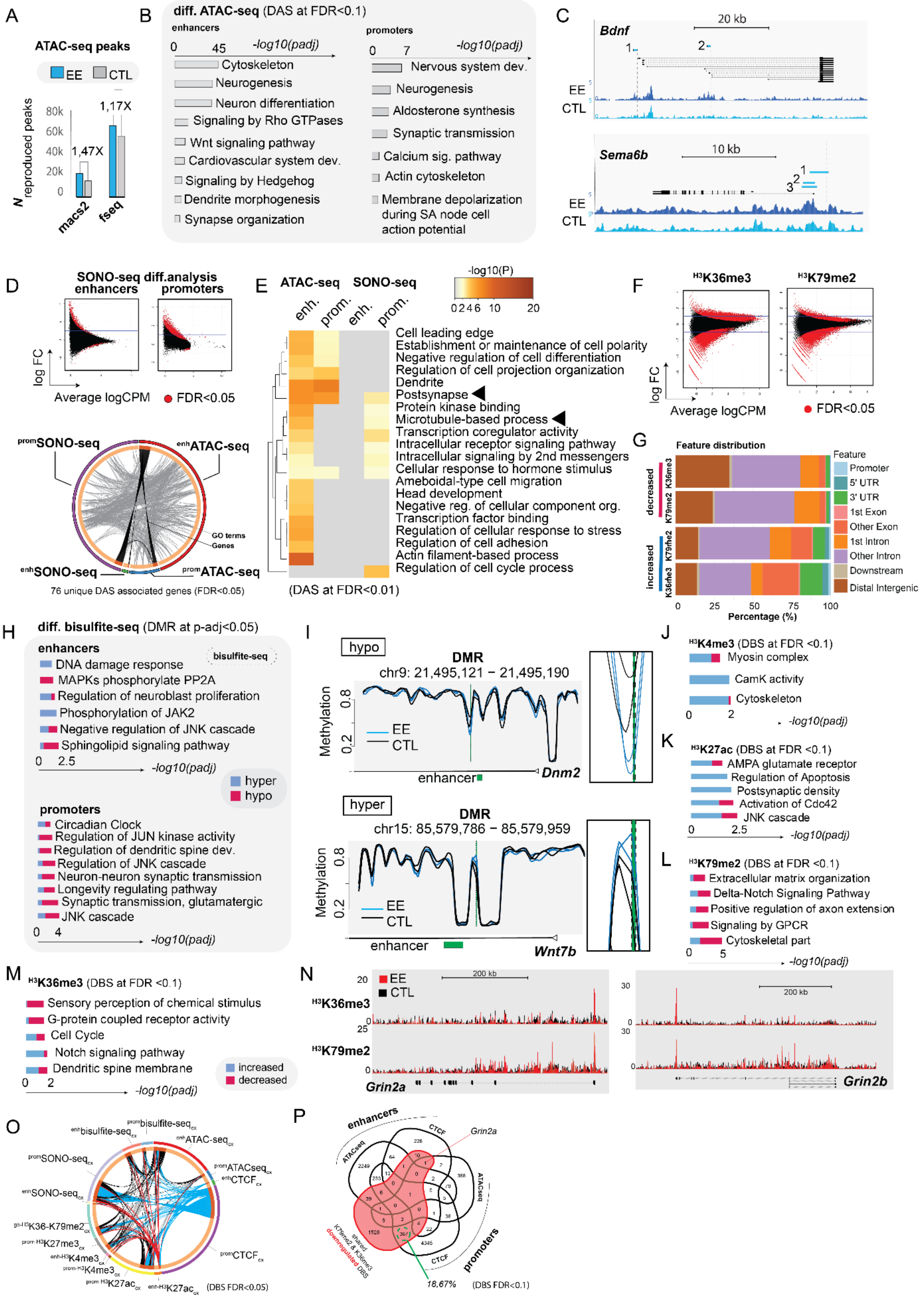
A
ATAC-seq peaks
EE
CTL
N reproduced peaks
80k
60k
40k
20k
0
1,47X
1,17X
macs2
fseq
B
diff. ATAC-seq (DAS at FDR<0.1)
enhancers
0
45
-log10(padj)
Cytoskeleton
Neurogenesis
Neuron differentiation
Signaling by Rho GTPases
Wnt signaling pathway
Cardiovascular system dev.
Signaling by Hedgehog
Dendrite morphogenesis
Synapse organization
promoters
0
7
-log10(padj)
Nervous system dev.
Neurogenesis
Aldosterone synthesis
Synaptic transmission
Calcium sig. pathway
Actin cytoskeleton
Membrane depolarization during SA node cell action potential
C
Bdnf
20 kb
EE
CTL
Sema6b
10 kb
EE
CTL
D
SONO-seq diff.analysis
enhancers
promoters
log FC
Average logCPM
FDR<0.05
promSONO-seq
enhATAC-seq
enhSONO-seq
promATAC-seq
GO terms
Genes
76 unique DAS associated genes (FDR<0.05)
E
ATAC-seq
SONO-seq
enh.
prom.
enh.
prom.
-log10(P)
0 2 4 6 10 20
Cell leading edge
Establishment or maintenance of cell polarity
Negative regulation of cell differentiation
Regulation of cell projection organization
Dendrite
Postsynapse
Protein kinase binding
Microtubule-based process
Transcription coregulator activity
Intracellular receptor signaling pathway
Intracellular signaling by 2nd messengers
Cellular response to hormone stimulus
Ameboidal-type cell migration
Head development
Negative reg. of cellular component org.
Transcription factor binding
Regulation of cellular response to stress
Regulation of cell adhesion
Actin filament-based process
Regulation of cell cycle process
(DAS at FDR<0.01)
F
H3K36me3
H3K79me2
log FC
Average logCPM
FDR<0.05
G
Feature distribution
decreased
increased
K36me3
K79me2
K79me2
K36me3
Feature
Promoter
5' UTR
3' UTR
1st Exon
Other Exon
1st Intron
Other Intron
Downstream
Distal Intergenic
0 25 50 75 100
Percentage (%)
H
diff. bisulfite-seq (DMR at p-adj<0.05)
enhancers
bisulfite-seq
DNA damage response
MAPKs phosphorylate PP2A
Regulation of neuroblast proliferation
Phosphorylation of JAK2
Negative regulation of JNK cascade
Sphingolipid signaling pathway
0 2.5
-log10(padj)
hyper
hypo
promoters
Circadian Clock
Regulation of JUN kinase activity
Regulation of dendritic spine dev.
Regulation of JNK cascade
Neuron-neuron synaptic transmission
Longevity regulating pathway
Synaptic transmission, glutamatergic
JNK cascade
0 4
-log10(padj)
I
hypo
DMR
chr9: 21,495,121 − 21,495,190
Methylation
0.8
0.2
EE
CTL
enhancer
Dnm2
hyper
DMR
chr15: 85,579,786 − 85,579,959
Methylation
0.8
0.2
EE
CTL
enhancer
Wnt7b
J
H3K4me3 (DBS at FDR <0.1)
Myosin complex
CamK activity
Cytoskeleton
0 2
-log10(padj)
K
H3K27ac (DBS at FDR <0.1)
AMPA glutamate receptor
Regulation of Apoptosis
Postsynaptic density
Activation of Cdc42
JNK cascade
0 2.5
-log10(padj)
L
H3K79me2 (DBS at FDR <0.1)
Extracellular matrix organization
Delta-Notch Signaling Pathway
Positive regulation of axon extension
Signaling by GPCR
Cytoskeletal part
0 5
-log10(padj)
M
H3K36me3 (DBS at FDR <0.1)
Sensory perception of chemical stimulus
G-protein coupled receptor activity
Cell Cycle
Notch signaling pathway
Dendritic spine membrane
0 2
-log10(padj)
increased
decreased
N
EE
CTL
200 kb
H3K36me3
H3K79me2
Grin2a
200 kb
Grin2b
O
enhbisulfite-seqcx
prombisulfite-seqcx
enhATAC-seqcx
promSONO-seqcx
promATACseqcx
enhCTCFcx
enhSONO-seqcx
gb-H3K36-K79me2cx
prom-H3K27me3cx
enh-H3K4me3cx
prom-H3K4me3cx
prom-H3K27accx
enh-H3K27accx
promCTCFcx
(DBS FDR<0.05)
P
enhancers
CTCF
Grin2a
ATACseq
ATACseq
promoters
CTCF
K79me2 & K36me3 shared downregulated DBS
18,67%
(DBS FDR<0.1)

**Figure S1. Chromatin accessibility and epigenetic changes induce by EE in cortex homogenate.** ATAC-seq, SONO-seq, bisulfite-seq and histone marks results at postnatal day [P51]. **A)** Enrichment of EE and CTL samples over a consensus peak-dataset computed using both macs2 and fseq. Bar-plot shows that the number of peaks found by macs2 and fseq was greater in EE compared to CTL samples. **B)** Gene ontology analysis of previous regulatory regions: enhancers (left bar-plot) and promoters (right) (Table S2, p-value < 0.05 – Bonferroni adjusted). **C)** Cortex homogenate UCSC screenshot examples of ATAC-seq differential accessibility sites at P51, showing increased accessibility in the promoter regions of *Bdnf* and *Sema6a* genes. Numbers 1 and 2 correspond to fseq/macs2 peak calls and number 3 to the defined promoter region (Method Details). **D)** MA-plot showing the log(fold change) vs log(counts per million) of differential SONO-seq signal on enhancers (left) and promoters (right) at P51 due EE (red dots are significant sites at FDR<0.05, Table S2, Method Details). Circos plot show the intersection of SONOseq and ATAC-seq differential results due EE at FDR<0.05. **E)** Metascape gene ontology analysis of differential enhancer associated genes and promoters in both ATAC-seq and SONO-seq at FDR<0.01. **F)** Differential analysis of H3K36me3 (left MA-plot) and H3K79me2 (right) on enhancer regions (red dots denote significant results at FDR<0.05. **G)** Genome distribution plot of enhancers showing differential activity of H3K36me3 and H3K79me2 due EE. Around 15-25% of these regions are distal intergenic enhancers. **H**) GO analysis (at p-value < 0.05 Benjamini-Hochberg adjusted) of bisulfite sequencing differential analysis of enhancers (upper bar-plot) and promoters (bellow). **I)** Selected examples of hypo (upper plot) and hyper methylation regions (bellow). **J, K, L, M)** GO analysis of H3K4me3 (K), H3K27ac (L), H3K79me2 (M) and H3K36me3 (N) p-value < 0.05 Benjamini-Hochberg adjusted**. N)** UCSC genome browser examples of glutamatergic genes showing decreased binding of H3K79me2 and H3K36me3 upon EE. **O)** Circos-plot showing the intersection of differential analysis in all cortical epigenetic marks upon EE at FDR<0.05. Highlighted in blue differential CTCF binding and in red H3K27ac. **P)** Venn-plot of potential impact into transcription-associated marks due differential activity shown in regulatory regions: enhancers and promoters. Associated-genes of ATAC-seq and CTCF showing corresponding differential enhancer and promoter activity intersected with consensus differential H3K79me2 & H3K36me regions (i.e. genes that shared differential gene-body activity due EE in both histone marks).

### Figure S2

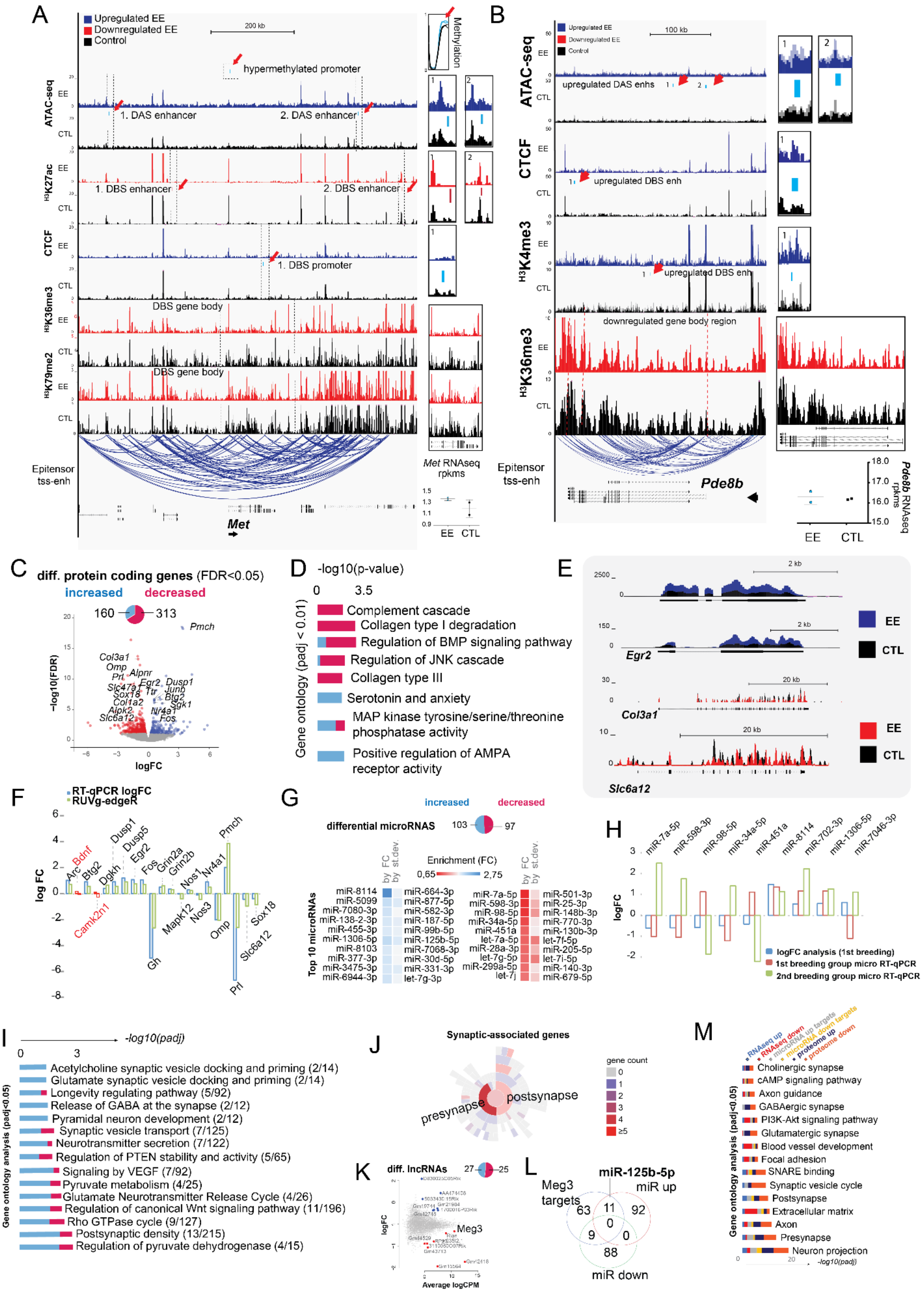
A
Upregulated EE
Downregulated EE
Control
hypermethylated promoter
Methylation
ATAC-seq
1. DAS enhancer
2. DAS enhancer
H3K27ac
1. DBS enhancer
2. DBS enhancer
CTCF
1. DBS promoter
H3K36me3
DBS gene body
H3K79me2
DBS gene body
Epitensor tss-enh
Met
Met RNAseq rpkms
EE
CTL
B
upregulated DAS enhs
upregulated DBS enh
H3K4me3
upregulated DBS enh
downregulated gene body region
Pde8b
Pde8b RNAseq rpkms
C
diff. protein coding genes (FDR<0.05)
increased
decreased
160
313
Pmch
-log10(FDR)
logFC
D
-log10(p-value)
Gene ontology (padj < 0.01)
Complement cascade
Collagen type I degradation
Regulation of BMP signaling pathway
Regulation of JNK cascade
Collagen type III
Serotonin and anxiety
MAP kinase tyrosine/serine/threonine phosphatase activity
Positive regulation of AMPA receptor activity
E
Egr2
Col3a1
Slc6a12
F
RT-qPCR logFC
RUVg-edgeR
log FC
G
differential microRNAs
103
97
Enrichment (FC)
Top 10 microRNAs
H
logFC
logFC analysis (1st breeding)
1st breeding group micro RT-qPCR
2nd breeding group micro RT-qPCR
I
-log10(padj)
Gene ontology analysis (padj<0.05)
Acetylcholine synaptic vesicle docking and priming (2/14)
Glutamate synaptic vesicle docking and priming (2/14)
Longevity regulating pathway (5/92)
Release of GABA at the synapse (2/12)
Pyramidal neuron development (2/12)
Synaptic vesicle transport (7/125)
Neurotransmitter secretion (7/122)
Regulation of PTEN stability and activity (5/65)
Signaling by VEGF (7/92)
Pyruvate metabolism (4/25)
Glutamate Neurotransmitter Release Cycle (4/26)
Regulation of canonical Wnt signaling pathway (11/196)
Rho GTPase cycle (9/127)
Postsynaptic density (13/215)
Regulation of pyruvate dehydrogenase (4/15)
J
Synaptic-associated genes
gene count
presynapse
postsynapse
K
diff. lncRNAs
27
25
Meg3
logFC
Average logCPM
L
miR-125b-5p
Meg3 targets
miR up
miR down
M
RNAseq up
RNAseq down
microRNA up targets
microRNA down targets
proteome up
proteome down
Cholinergic synapse
cAMP signaling pathway
Axon guidance
GABAergic synapse
PI3K-Akt signaling pathway
Glutamatergic synapse
Blood vessel development
Focal adhesion
SNARE binding
Synaptic vesicle cycle
Postsynapse
Extracellular matrix
Axon
Presynapse
Neuron projection
-log10(padj)
Gene ontology analysis (padj<0.05)

**Figure S2. Transcriptional and translational changes due EE. A, B)** *Met* and *Pde8b* USCS screenshot examples showing gene-body downregulation of H3K36me3 and H3K79me2 together with differential CTCF and ATAC-seq signals upon EE**. C)** Differential expression of poly-A expression volcano plot representing changes found using RUV-g correction method. Blue, upregulated genes; red, downregulated genes at FDR<0.05. **D)** Gene ontology analysis of selected significant (p-value < 0.01 Bejamini-Hochberg corrected) terms of differential expressed genes (DEG). Blue bars represent the percentage of the p-value enrichment of genes that are upregulated and pink the proportion of downregulated genes. **E)** Screenshot of genome browser DEG examples. **F)** poly-A validation plot by qPCR. Blue bars represent the logFC of EE vs CTL measured by qPCR and compared to DEG found in the differential analysis using RUV-g correction. **G)** Top differential upregulated (left) and downregulated (right) microRNAs sorted by fold change and p-value. Up-right, circle-plot of total number of differential microRNAs found in our data. **H)** Validation of microRNAs in different batch set of animals by qPCR (Method Details). **I)** microRNA targets gene ontology (p-adj <0.05). **J)** SynGo synapse enrichment analysis of microRNA targets. **K)** Differential expression of lncRNAs highlighting *Meg3.* **L)** Intersection of *Meg3* predicted microRNA binding with LncBase v2.0 with differential microRNAs found in our dataset (Table S4). **M)** Gene ontology analysis of significant terms (p-value < 0.05 adjusted) found in the differential analysis of RNAseq, microRNA and proteome.

### Figure S3

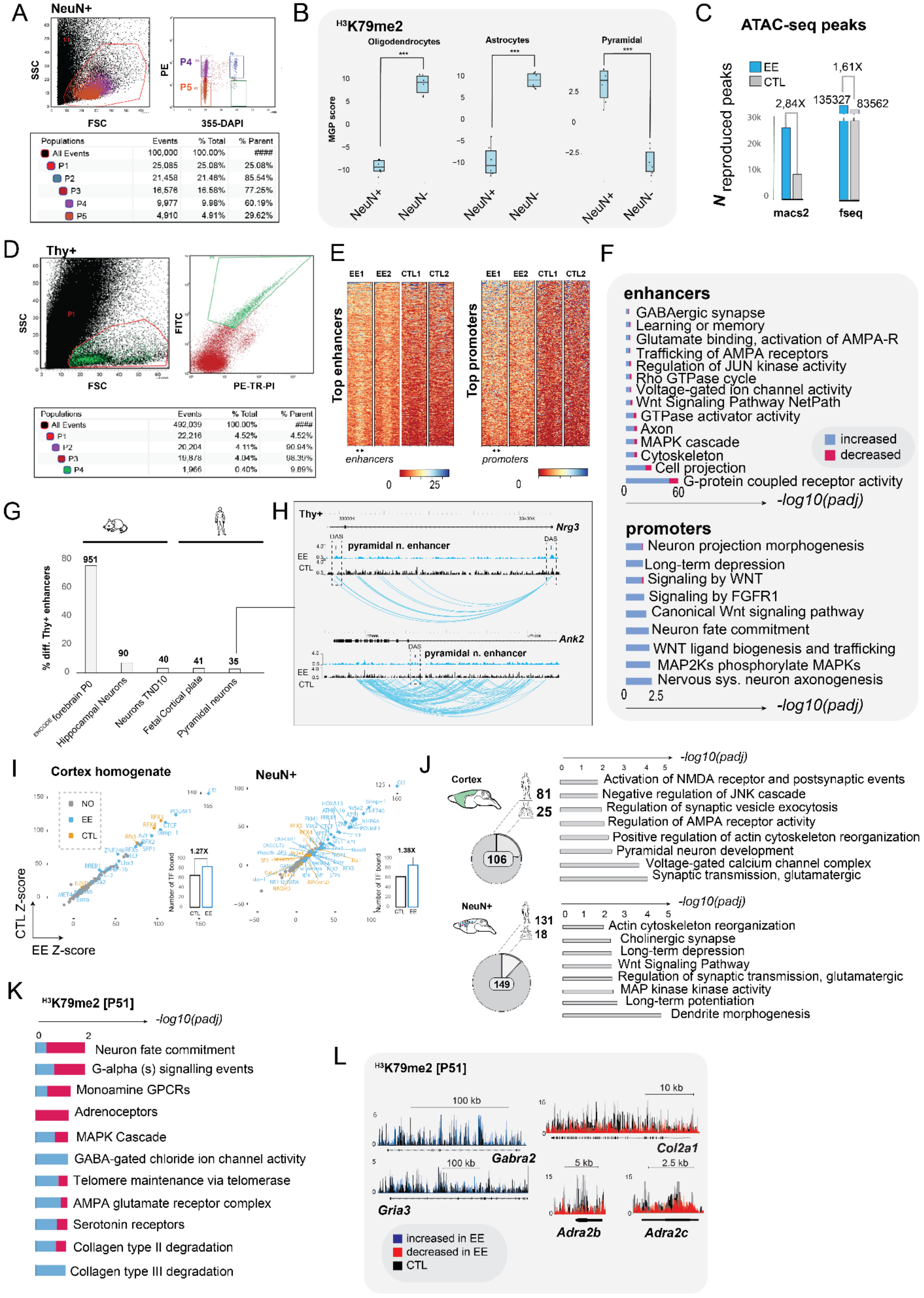
A
NeuN+
SSC
FSC
PE
P4
P5
355-DAPI
Populations Events % Total % Parent
All Events 100,000 100.00% ####
P1 25,085 25.08% 25.08%
P2 21,458 21.46% 85.54%
P3 16,576 16.58% 77.25%
P4 9,977 9.98% 60.19%
P5 4,910 4.91% 29.62%
B
H3K79me2
Oligodendrocytes
Astrocytes
Pyramidal
MGP score
NeuN+
NeuN-
C
ATAC-seq peaks
EE
CTL
1,61X
2,84X
135327
83562
N reproduced peaks
macs2
fseq
D
Thy+
SSC
FSC
FITC
PE-TR-PI
Populations Events % Total % Parent
All Events 492,039 100.00% ####
P1 22,216 4.52% 4.52%
P2 20,204 4.11% 90.94%
P3 19,878 4.04% 98.39%
P4 1,966 0.40% 9.89%
E
EE1 EE2 CTL1 CTL2
Top enhancers
enhancers
Top promoters
promoters
F
enhancers
GABAergic synapse
Learning or memory
Glutamate binding, activation of AMPA-R
Trafficking of AMPA receptors
Regulation of JUN kinase activity
Rho GTPase cycle
Voltage-gated ion channel activity
Wnt Signaling Pathway NetPath
GTPase activator activity
Axon
MAPK cascade
Cytoskeleton
Cell projection
G-protein coupled receptor activity
increased
decreased
-log10(padj)
promoters
Neuron projection morphogenesis
Long-term depression
Signaling by WNT
Signaling by FGFR1
Canonical Wnt signaling pathway
Neuron fate commitment
WNT ligand biogenesis and trafficking
MAP2Ks phosphorylate MAPKs
Nervous sys. neuron axonogenesis
-log10(padj)
G
% diff. Thy+ enhancers
951
90
40
41
35
ENCODE forebrain P0
Hippocampal Neurons
Neurons TND10
Fetal Cortical plate
Pyramidal neurons
H
Thy+
Nrg3
pyramidal n. enhancer
DAS
EE
CTL
Ank2
I
Cortex homogenate
NeuN+
NO
EE
CTL
1.27X
1.38X
Number of TF bound
CTL Z-score
EE Z-score
J
Cortex
81
25
106
-log10(padj)
Activation of NMDA receptor and postsynaptic events
Negative regulation of JNK cascade
Regulation of synaptic vesicle exocytosis
Regulation of AMPA receptor activity
Positive regulation of actin cytoskeleton reorganization
Pyramidal neuron development
Voltage-gated calcium channel complex
Synaptic transmission, glutamatergic
NeuN+
131
18
149
-log10(padj)
Actin cytoskeleton reorganization
Cholinergic synapse
Long-term depression
Wnt Signaling Pathway
Regulation of synaptic transmission, glutamatergic
MAP kinase kinase activity
Long-term potentiation
Dendrite morphogenesis
K
H3K79me2 [P51]
-log10(padj)
Neuron fate commitment
G-alpha (s) signalling events
Monoamine GPCRs
Adrenoceptors
MAPK Cascade
GABA-gated chloride ion channel activity
Telomere maintenance via telomerase
AMPA glutamate receptor complex
Serotonin receptors
Collagen type II degradation
Collagen type III degradation
L
H3K79me2 [P51]
100 kb
Gabra2
100 kb
Gria3
10 kb
Col2a1
5 kb
Adra2b
2.5 kb
Adra2c
increased in EE
decreased in EE
CTL

**Figure S3. Chromatin accessibility and transcription-associated changes due EE in sorted populations. A)** Gating strategy: FAC-sorting report plots of extracted NeuN+ and NeuN- populations. Plot y-axis shows side-scattered light (SSC) versus x-axis Forward-scattered light (FSC) plot of NeuN+ nuclei preparation. The SSC value reports the complexity of the sample measured by the refraction of the laser light beamed in the surface of the nuclei or cells in the flow. The FSC reports the intensity of the signal given by the conjugated antibody (Alexa Fluor-555). Right**,** intensity plot of the signal of the conjugated antibody (PE, Alexa-555) versus DAPI intensity. P4 and P6 represent the nuclei positive for the neural marker NeuN+. **B)** Cell specificity of H3K79me2 NeuN+ and NeuN- populations using curated gene specific cell markers discovered by single-cell RNAseq (Mancarci et al., 2017) (Method Details). Neuronal and non-neuronal H3K79me2 counts in gene body regions were normalized to rpkms. The cell specify is assessed by the marker gene profile (MGP) score, and it was calculated for oligodendrocytes (left), astrocytes (center) and pyramidal neurons (right) markers. Significant oligodendrocytes and astrocytes (pvalue< 0.01) markers were found to be associated more to NeuN- populations, meanwhile significant pyramidal markers (pvalue<0.01) were found to be associated to NeuN+.**C)** Enrichment of EE and CTL samples over a consensus peak-dataset computed using both macs2 and fseq in NeuN+ sorted nuclei. Bar-plot shows that the number of peaks found by macs2 and fseq was greater in EE compared to CTL samples. **D)** Side-scattered light (SSC) versus Forward-scattered ligh (FSC) plot of YFP+ pyramidal neurons preparation (Tg-Thy1+mice, left), and FITC signal of the YFP neuronal cells (right). **E)** Top changes of ATAC-seq in regulatory regions of Thy1+ pyramidal neurons (n=2 per condition). **F)** Gene ontology analysis of the Thy+ differential accessibility sites in enhancers (upper plot) and promoters (bellow). **G)** Thy+ enhancer intersection showing differential accessibility due EE with mouse ENCODE forebrain P0 enhancers (ENCFF711DUZ), hippocampal and TND10 mature neurons and lifted mouse to human intersection with human fetal cortical plate and pyramidal neuron enhancers (Dong et al., 2018; Fernandez-Albert et al., 2019; de la Torre-Ubieta et al., 2018; Thakurela et al., 2015) **H)** Epigenome WashU screenshot of shared enhancers with human pyramidal neurons lifted to mouse genomic coordinates. **I)** Footprint analysis. Z-scores regression plots showing transcription factors binding (control - orange, EE - blue), bar-plots represent the accumulated number of TF bound in both conditions in whole cortex (left) and NeuN+ (right). **J)** CTCF bound genes found for cortex (upper plot) and NeuN+ samples (bellow). Motifs detected were human and *Drosophila* specific. Drawings (up-right of the pie-plot) assigned the number instances found per specie. Next to it (right) associated gene ontology analysis of selected terms (pvalue < 0.05 Benjamini-Holchberg adjusted, Table S2). **K)** H3K79me2 gene ontology analysis of EE vs CTL at P51 (p-value < 0.05, Benjamini-Hochberg adjusted) **L**) NeuN+ H3K79me3 at P51 UCSC screenshots of increased (blue) and decreased (red) binding due EE stimulation versus CTL samples (black tracks).

#### Figure S4

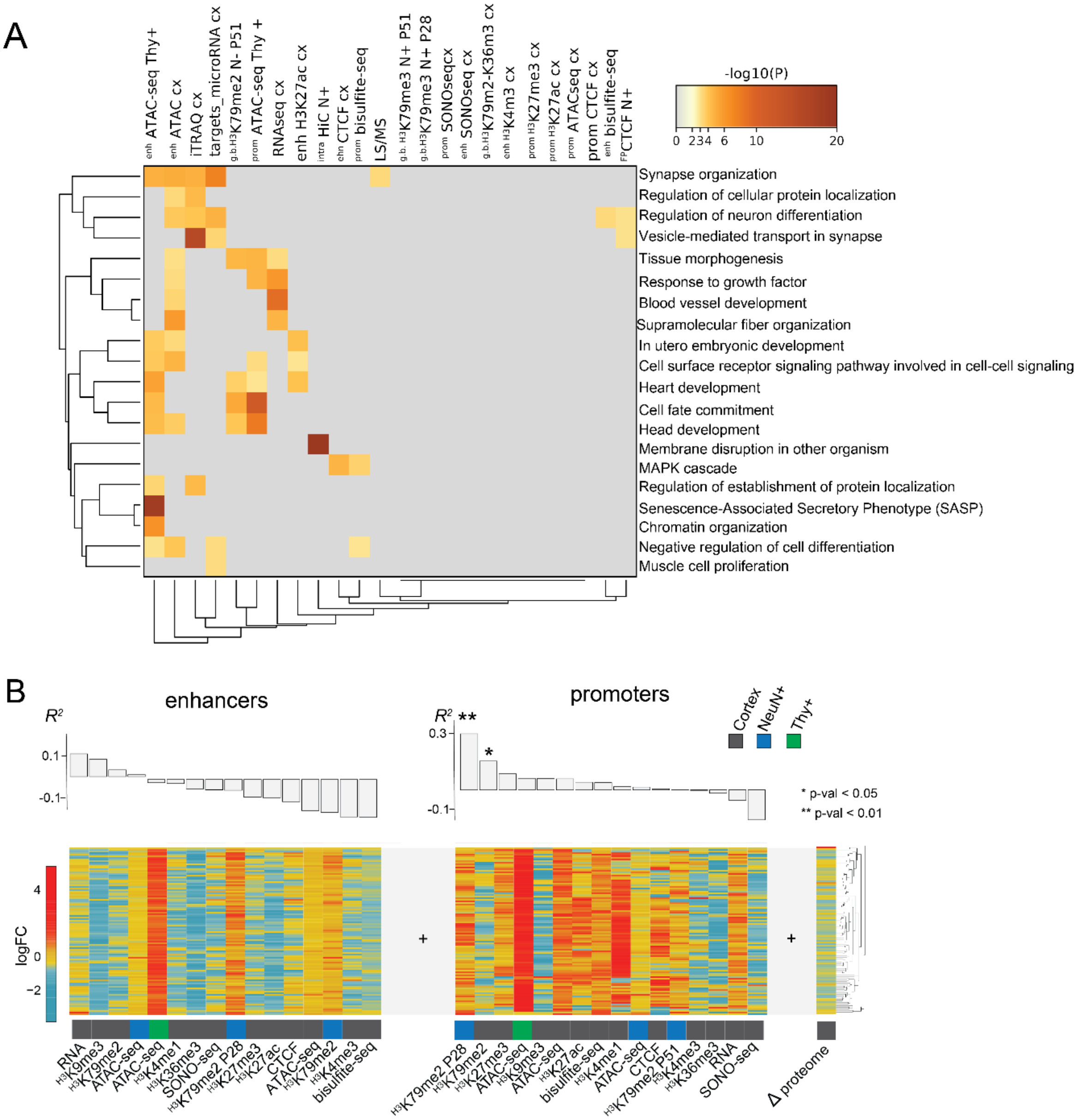
A
-log10(P)
0 2 3 4 6 10 20
Synapse organization
Regulation of cellular protein localization
Regulation of neuron differentiation
Vesicle-mediated transport in synapse
Tissue morphogenesis
Response to growth factor
Blood vessel development
Supramolecular fiber organization
In utero embryonic development
Cell surface receptor signaling pathway involved in cell-cell signaling
Heart development
Cell fate commitment
Head development
Membrane disruption in other organism
MAPK cascade
Regulation of establishment of protein localization
Senescence-Associated Secretory Phenotype (SASP)
Chromatin organization
Negative regulation of cell differentiation
Muscle cell proliferation
B
enhancers
promoters
Cortex
NeuN+
Thy+
* p-val < 0.05
** p-val < 0.01
logFC
Δ proteome

**Figure S4. Data integration and regulatory regions activity translated into proteomic changes. A)** Metascape gene ontology analysis of all differential changes induced by EE at FDR<0.05. **B)** Spearman translation efficiency of epigenetic marks in enhancers and promoters into differentially proteomic changes.
